# Advection versus diffusion in brain ventricular transport

**DOI:** 10.1101/2025.04.15.648743

**Authors:** Halvor Herlyng, Ada J. Ellingsrud, Miroslav Kuchta, Inyoung Jeong, Marie E. Rognes, Nathalie Jurisch-Yaksi

**Affiliations:** Department of Numerical Analysis and Scientific Computing, Simula Research Laboratory, Oslo, Norway; Department of Clinical and Molecular Medicine, Norwegian University of Science and Technology, Trondheim, Norway; K. G. Jebsen Center for Brain Fluid Research, Oslo, Norway

**Keywords:** cerebrospinal fluid, brain ventricles, cilia, transport, finite elements

## Abstract

Cerebrospinal fluid (CSF) is integral to brain function. CSF provides mechanical support for the brain and helps distribute nutrients, neurotransmitters and metabolites throughout the central nervous system. CSF flow is driven by several processes, including the beating of motile cilia located on the walls of the brain ventricles. Despite the physiological importance of CSF, the underlying mechanisms of CSF flow and solute transport in the brain ventricles remain to be comprehensively resolved. This study analyzes and evaluates specifically the role of motile cilia in CSF flow and transport. We developed finite element methods for modeling flow and transport using the geometry of the zebrafish larval brain ventricles, for which we have detailed knowledge of cilia properties and CSF motion. The computational model is validated by in vivo experiments that monitor transport of a photoconvertible protein secreted in the brain ventricles. Our results show that while cilia contribute to advection of large particles, diffusion plays a significant role in the transport of small solutes. We also demonstrate how cilia location and the geometry of the ventricular system impact solute distribution. Altogether, this work presents a computational framework that can be applied to other ventricular systems, together with new concepts of how molecules are transported within the brain and its ventricles.

## 1 Introduction

Brain development and function depend on numerous regulatory processes, including the flow of cerebrospinal fluid (CSF).^[1]^ Produced predominantly by the choroid plexus, CSF is a clear liquid that flows through the ventricular system and subarachnoid space.^[2–4]^ By transporting growth factors, nutrients, neurotransmitters and metabolites, as well as removing waste products, CSF regulates brain development and homeostasis.^[1, 5, 6]^ Disruption of CSF circulation is detrimental to brain health and disturbs cognitive and motor functions,^[7]^ and accumulation of CSF leads to hydrocephalus, a disorder often requiring emergency surgical treatment.^[8–11]^

The flow of CSF in the ventricular system is driven by multiple physiological processes, including the beating motion of motile cilia, CSF secretion by the choroid plexus, and pulsatile pressure gradients generated by the cardiac cycle, respiration, neural activity, and body movements.^[2, 5, 6, 12–17]^ Cilia are hair-like structures found in a vast diversity of biological organisms and in several human organs, including the brain.^[5, 18–29]^ Notably, motile cilia are found on the ventricular surface of the brain, known as the ependyma, and beat in a coordinated pattern at frequencies of 10–40 Hz to mediate fluid flow.^[5, 23, 25, 30, 31]^ Given their microscopic size, ranging from 5 to 15 microns, cilia contribute primarily to flow near the ependyma in the brain.^[5, 14, 32, 33]^ In contrast, cardiac, respiratory and delta waves at lower frequencies induce pulsatile flow at larger scales in the brain ventricular system, subarachnoid space, and perivascular spaces.^[12, 16, 34–37]^

To date, it remains poorly understood how motile cilia contribute to the overall fluid flow and solute transport within the brain. Experimental approaches face technical challenges: the microscopic size and relatively high beating frequencies of the cilia,^[14, 23, 38–41]^ as compared to other slower fluid dynamic processes that span larger spatial scales,^[12, 16, 34, 36]^ make cilia difficult to monitor in vivo in the rodent and human brain. Until now, most studies have analyzed one physical mechanism at a time, especially in mammals. For instance, studies using brain explants have convincingly shown that cilia generate complex flow patterns at the surface of the brain ventricles;^[33, 42, 43]^ however, they did not study the role of other contributions to CSF flow and the accompanying transport, such as pressure gradients associated with respiration and cardiac pulsations. Moreover, while MRI technology can measure CSF motion in humans,^[44–46]^ it cannot detect the microscopic movements of cilia to understand their impact in the large human brain.^[32]^ These technical limitations of studying cilia-mediated flow and transport make mathematical modeling and computational investigations an attractive complement to experimental studies. Previous simulation studies have focused on modeling the interaction between single or several cilia and surrounding fluids.^[47–51]^ Others considered cilia-mediated flow in the brain,^[29, 32, 52, 53]^ and other applications such as ciliary transport of mucous and particles in airways,^[19, 31, 54]^ propulsion and locomotion of ciliated organisms,^[21, 22, 27]^ and oxygen regulation in coral reefs.^[20]^ These simulation studies have provided valuable insights on how cilia mediate complex biological flows, and how cilia may contribute to tissue physiology. However, we still lack a comprehensive understanding of how cilia contribute to CSF motion and solute transport within the brain. This is of particular importance since disorders such as hydrocephalus are observed in animal models and humans with deficient motile cilia,^[14, 55–57]^ indicating that the cilia play a crucial role for healthy brain function.

In this study, we combined computational and experimental approaches to investigate the role of motile cilia in brain ventricular CSF flow and transport. We model the flow and transport by partial differential equations and solve them numerically with finite element methods to compute flow patterns and solute concentration dynamics. A geometry representing the zebrafish larval brain ventricles was used, for which we have detailed knowledge of cilia properties and CSF motion velocities and frequencies.^[14]^ Notably, this early developmental stage allows us to simplify our model by omitting CSF secretion, which starts at later stages,^[58]^ while still being well-conserved with mammals.^[5, 14, 18, 41, 59]^ The computational model is validated against experimental data by monitoring in vivo the dynamics of a photoconvertible fluorescent protein Dendra2 expressed within the brain ventricles. Our results show that diffusion plays a major role in the transport of small solutes, whereas cilia are paramount for the movement of larger particles. We also found that the location of cilia, as well as changes to the ventricular geometry, largely impact solute distribution. In summary, our work presents a computational framework that can be applied to other ventricular systems in the future, together with new concepts of how cilia distribute molecules and particles within the brain ventricles.

## 2 Results

### 2.1 Motile cilia induce flow compartmentalization within the brain ventricles

In vivo particle tracking reveals characteristic CSF flow patterns in zebrafish larval brain ventricles (Figure 1a– c).^[14]^ We attempt to accurately represent these flow patterns with computational fluid dynamics driven by cardiac pulsations and motile cilia lining the surface of the brain ventricles.

**Figure 1:**
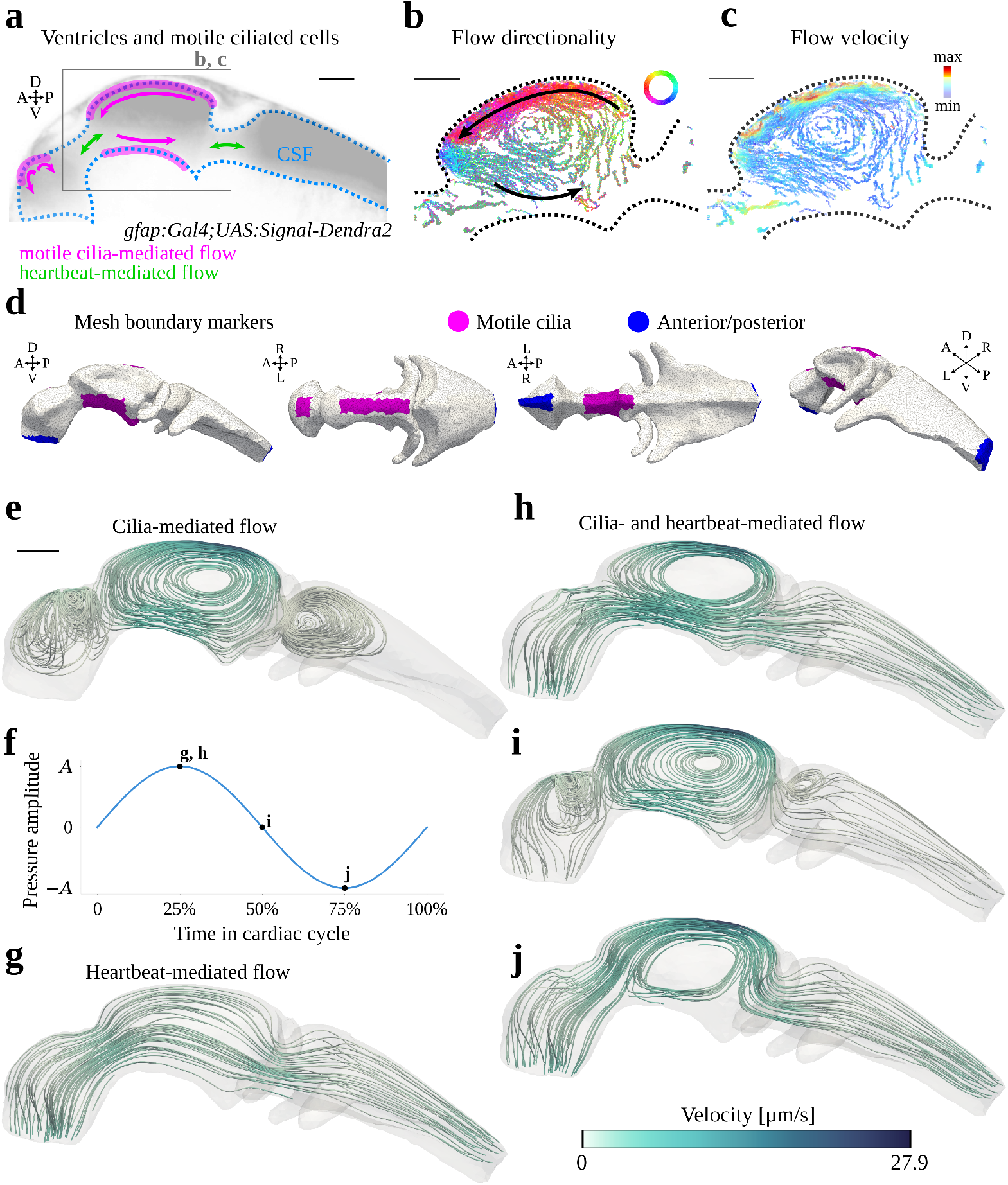
**a**. Schematic illustration of the two cerebrospinal fluid (CSF) flow components modeled, motile cilia and cardiac pulsations, and their contributions to the flow patterns. **b, c**. Particle tracking data visualizing CSF flow directionality (**b**) and velocity magnitudes (**c**) in the middle ventricle. **d**. Computational mesh with marked facets indicating regions of motile cilia (magenta) and anterior/posterior facets (blue). Dorsal (D), ventral (V), anterior (A), posterior (P), left (L) and right (R) coordinates indicated. **e**. Streamlines of the (steady-state) CSF flow field simulated with the cilia-driven/no-cardiac flow model. **f**. Illustration of the cardiac-related normal pressure forcing imposed on the anterior/posterior boundaries. Time instants of streamline plots indicated. **g**. Streamlines of the CSF flow field simulated with the cardiac-induced/no-cilia flow model at a time 25% into the cardiac cycle. **h–j**. Streamlines simulated with the baseline flow model (cilia+cardiac) at a time 25% (**h**), 50% (**i**) and 75% (**j**) into the cardiac cycle. Scale bars 50 µm.

Assuming that viscous forces are dominating the ventricular CSF flow, we model the flow by the Stokes equations. The net forces of motile cilia acting on the CSF are modeled as a steady, bidirectional tangential traction acting on parts of the ventricular wall (Figure 1d), with an impermeability condition (no normal flow) at the walls. Simulations show that this traction alone leads to a partial compartmentalization of the ventricular system, with large-scale vortex structures in the anterior, middle and posterior ventricles (Figure 1e). The highest flow speeds (26.7 µm/s) occur in the vicinity of the dorsal cilia in the middle ventricle. To model the cardiac pulsations, we impose a pulsatile pressure difference between the anterior and posterior boundaries (Figure 1d, f). The resulting flow is pulsatile and directional (Figure 1g), and in the absence of the cilia, the vortex structures vanish. The velocity magnitude peaks at 4.8 µm/s. CSF initially flows rostrocaudally before changing direction to caudorostral in the middle of the cardiac cycle.

When combining cilia motion and cardiac pulsations, we recognize flow features persisting throughout the cardiac cycle, now with vortex structures in the anterior and middle ventricles and directional flow in the posterior ventricle (Figure 1h–j, Supplementary Video S1). These characteristic patterns suggest that the flow in the anterior and middle ventricles is primarily driven by the motile cilia, while the cardiac pressure pulsations dominate within the posterior ventricle and in the ducts connecting the ventricular compartments. The CSF flow speed peaks at the time of peak caudal cardiac pulsatile flow, when both the tangential traction induced by the cilia and the pulsatile pressure-driven flow are aligned with the rostrocaudal axis, reaching up to 27.9 µm/s dorsally in the posterior part of the middle ventricle. The flow Reynolds number is Re ≈ 0.004, justifying the assumption of Stokes flow. In summary, the computational CSF flow model reproduces the flow features observed in vivo.^[14]^

### 2.2 Ventricular solute transport is advection-dominated for larger molecules

Proteins, neurotransmitters and other molecules intrinsically diffuse within the brain ventricles, but are also advected by the flow of CSF. A natural question is which of these mechanisms dominates for any given type of solute. To address this question, we simulate the distribution and evolution of three different solutes injected within a region of interest (ROI 1) in the dorsal region of the middle ventricle (Figure 2a) and subject to the CSF flow induced by motile cilia and cardiac pressure pulsations. In particular, we consider three different molecular diffusion coefficients, corresponding to a large extracellular vesicle (EV, *D*_1_ = 2.17 × 10^−12^ m^2^/s), a 90 kDa Starmaker-Green Fluorescent Protein (STM-GFP, *D*_2_ = 5.75 × 10^−11^ m^2^/s), and a 26 kDa photoconvertible protein Dendra2 (*D*_3_ = 1.15 × 10^−10^ m^2^/s).

**Figure 2:**
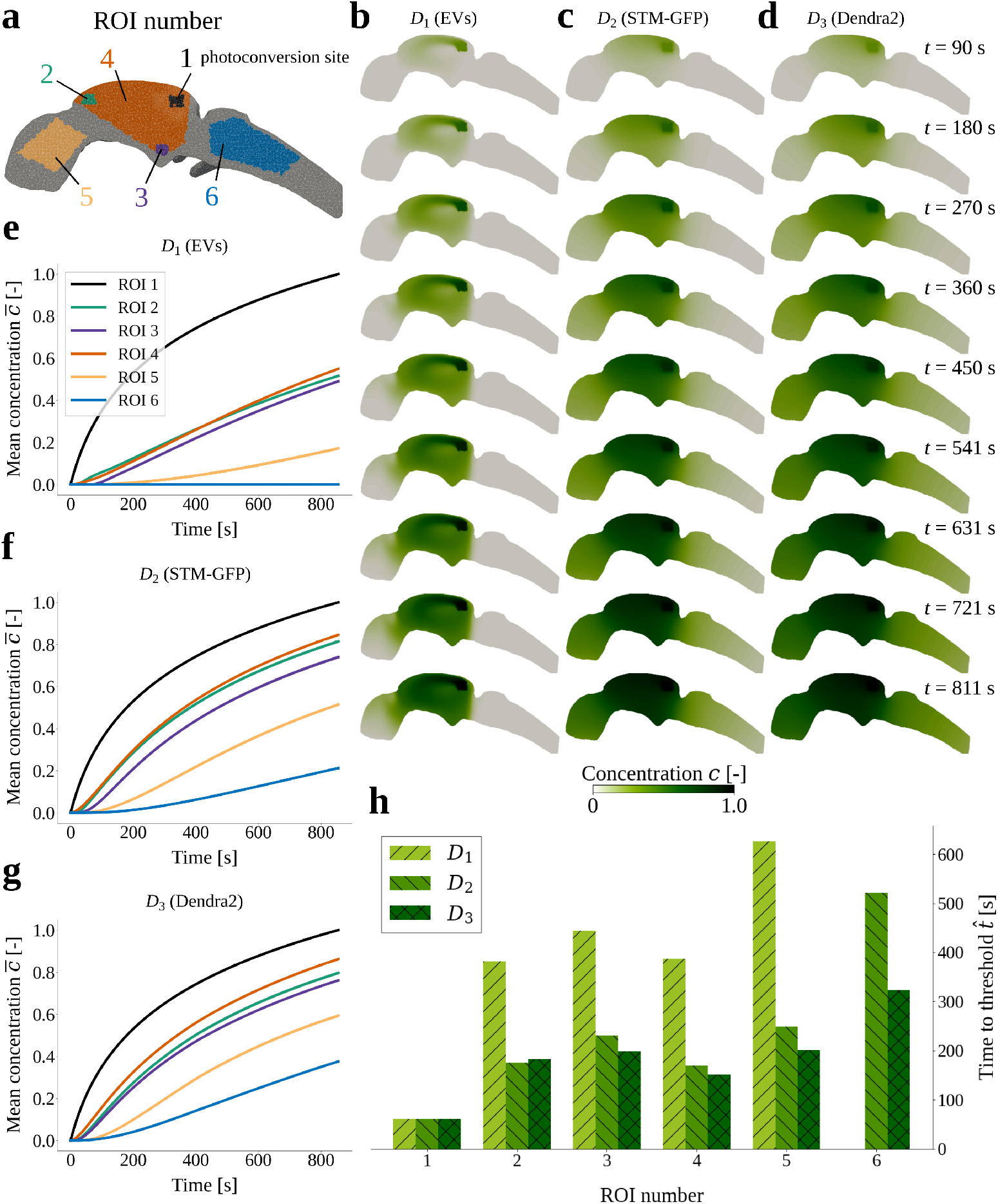
Simulated distribution and evolution after photoconversion of extracellular vesicles (EVs, *D*_1_), STM-GFP (*D*_2_) and Dendra2 (*D*_3_). **a**. Geometry-centered clip of the ventricles mesh showing the locations of the regions of interest (ROIs). **b–d**. Slices of the concentration field simulated with diffusion coefficients *D*_1_, *D*_2_ and *D*_3_ for nine time instants covering the time range from 90 seconds to 811 seconds, corresponding to 200 to 1800 cardiac cycles, for even intervals of 200 cycles. Slices are taken in the *xz*-plane (*y* = 0.140 mm, the center of the geometry). **e–g**. Mean concentration 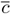 in each ROI as function of time for diffusion coefficients *D*_1_, *D*_2_ and *D*_3_. **h**. The time 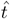 before the mean concentration 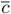 exceeds a threshold value of 0.25 (ROIs 2-4) or 0.10 (ROIs 5, 6) in each simulation setup. Note that there is no bar for *D*_1_ in ROI 6, because the threshold value was never exceeded.

Interestingly, the distribution patterns differ between the solutes. For the smallest diffusion coefficient, representing extracellular vesicles, the advective transport by the CSF dominates diffusion: the CSF flow structures are evident in the concentration field, indicating that transport along the streamlines is more rapid than the diffusion across the streamlines (Figure 2b). The balance between diffusion and advection shifts with increasing diffusion coefficient as solute spreads more uniformly throughout the ventricular geometry (Figure 2b–d). Notably, the smaller solutes with higher diffusion coefficients spread more quickly to distal regions. Computing Péclet numbers, we find Pe_1_ = 664, Pe_2_ = 25 and Pe_3_ = 13 for *D*_1_, *D*_2_ and *D*_3_, respectively. With all Pe *>* 1, this indicates that advection is the dominant transport mechanism on a global scale.

To further quantify the solute transport, we examine the mean concentrations 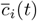 in six ROIs over time (Figure 2e–g) and, in addition, the times 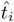 when the mean concentrations first exceed a threshold value *ĉ*_*i*_ (Figure 2h). Higher diffusion coefficients result in more rapid transport (Figure 2e–g), with lower times-to-threshold in ROIs 2–6 (Figure 2h). For all three diffusion coefficients, the solute first spreads within the middle ventricle, covering ROIs 2–4, before appearing in the anterior ventricle (ROI 5) and finally reaching the posterior ventricle (ROI 6). Notably, the extracellular vesicles, associated with the smallest diffusion coefficient, do not reach the posterior ventricle (ROI 6) within the simulated time frame Figure 2b, e.

### 2.3 Model validation by comparison with photoconversion experiments

To validate the model predictions, we performed photoconversion experiments in transgenic zebrafish larvae expressing the secreted Dendra2 in their brain ventricles. The approach is inspired by earlier work where a photoconvertible protein was injected into the ventricle.^[60]^ By using a transgenic system where the protein is directly secreted into the CSF, we avoid detrimental effects of intraventricular injection on CSF properties. Our photoconversion protocol consisted of the acquisition of a baseline fluorescence intensity *F*_0_, before localized exposure to a UV laser (at ROI 1) that converts Dendra2 from a green-to a red-emitting fluorescent protein (Supplementary Video S2). To quantify the transport of photoconverted proteins, we measured and calculated the change in fluorescence intensity Δ*F* (*t*) = (*F* (*t*) −*F*_0_)*/F*_0_ over time (Figure 3a).

**Figure 3:**
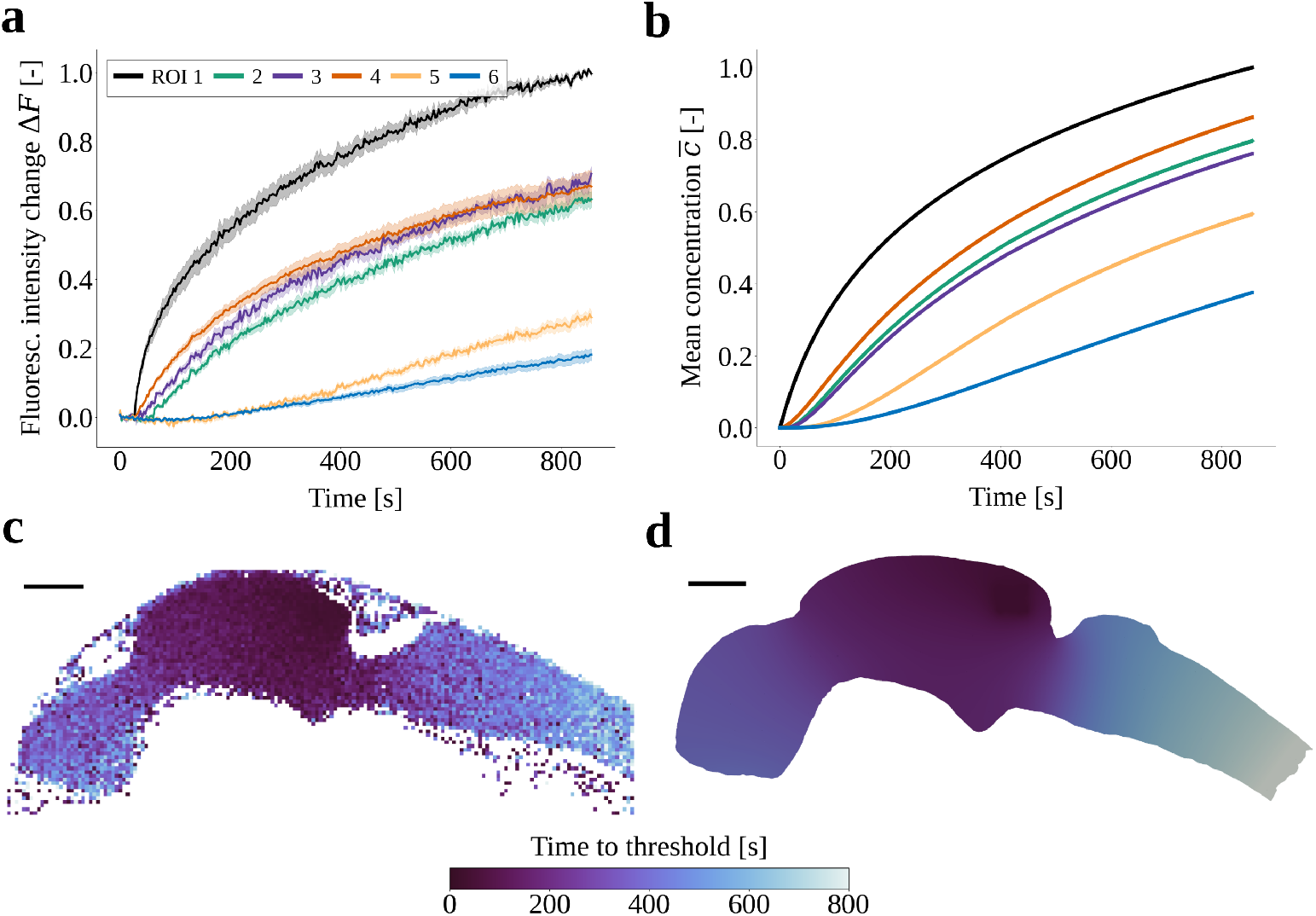
**a**. Relative change in fluorescence intensity Δ*F* (*t*) in each region of interest (ROI) observed during experiments monitoring the movement of Dendra2 proteins photoconverted by laser targeting ROI 1. Bold lines are mean values of the cohort (*n* = 8) and shaded regions cover one standard deviation. **b**. Mean concentration 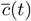 in each ROI in a simulation of Dendra2 transport with the diffusion coefficient *D*_3_. **c**. Time to Δ*F* (*t*) reaches a threshold value of 0.25 in one representative zebrafish larva. **d**. Time to 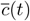 reaches a threshold value of 0.25 for a transport simulation with diffusion coefficient *D*_3_. The posterior-most regions of the geometries are excluded, because Δ*F* and 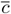 never exceeded the threshold value of 0.25 in these regions. Scale bars 50 µm.

Comparison of the intensity changes with the simulated concentration profiles in ROIs 1–6 (Figure 3a, b) shows that the model predictions agree well with the experimental measurements. We note minor differences however. The simulated values of 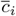 at the final time (0.80, 0.76, 0.86, 0.60 and 0.38 in ROIs 2–6, respectively) are generally higher than the corresponding experimental Δ*F* values (0.63, 0.71, 0.67, 0.29 and 0.18). Moreover, we observe differences in the distribution within the middle ventricle, especially with regard to ROIs 2 and 3. Overall, we note that the time to reach a 25% threshold agree well between the model predictions and experimental values (Figure 3c, d).

### 2.4 Absence of ciliary motion affects local solute distribution

Ciliary motion is integral to physiological CSF flow, and motile-cilia deficiency has been associated with defective brain development and various pathologies in zebrafish, rodents and humans.^[14, 33, 42, 55, 57, 61–66]^ As observed both experimentally and computationally, absence of motile cilia alters CSF flow patterns Figure 1g.^[14]^ We next investigate the implications of these changes on brain ventricular transport. We experimentally measured the movement of proteins by monitoring photoconverted Dendra2 signal in control zebrafish and motile-cilia mutant *schmalhans* (*smh, ccdc103*) zebrafish.^[67]^ In ROIs 2 and 4, which quantify Dendra2 distribution downstream of the photoconversion site, we observed a slower increase of fluorescence intensity (Figure 4a): the mean times-to-threshold in the *smh* mutants were 45% (ROI 2) and 25% (ROI 4) higher compared to controls (Figure 4b). We found no significant differences between *smh* mutants and controls in the Dendra2 distribution in ROIs 3, 5, and 6.

**Figure 4:**
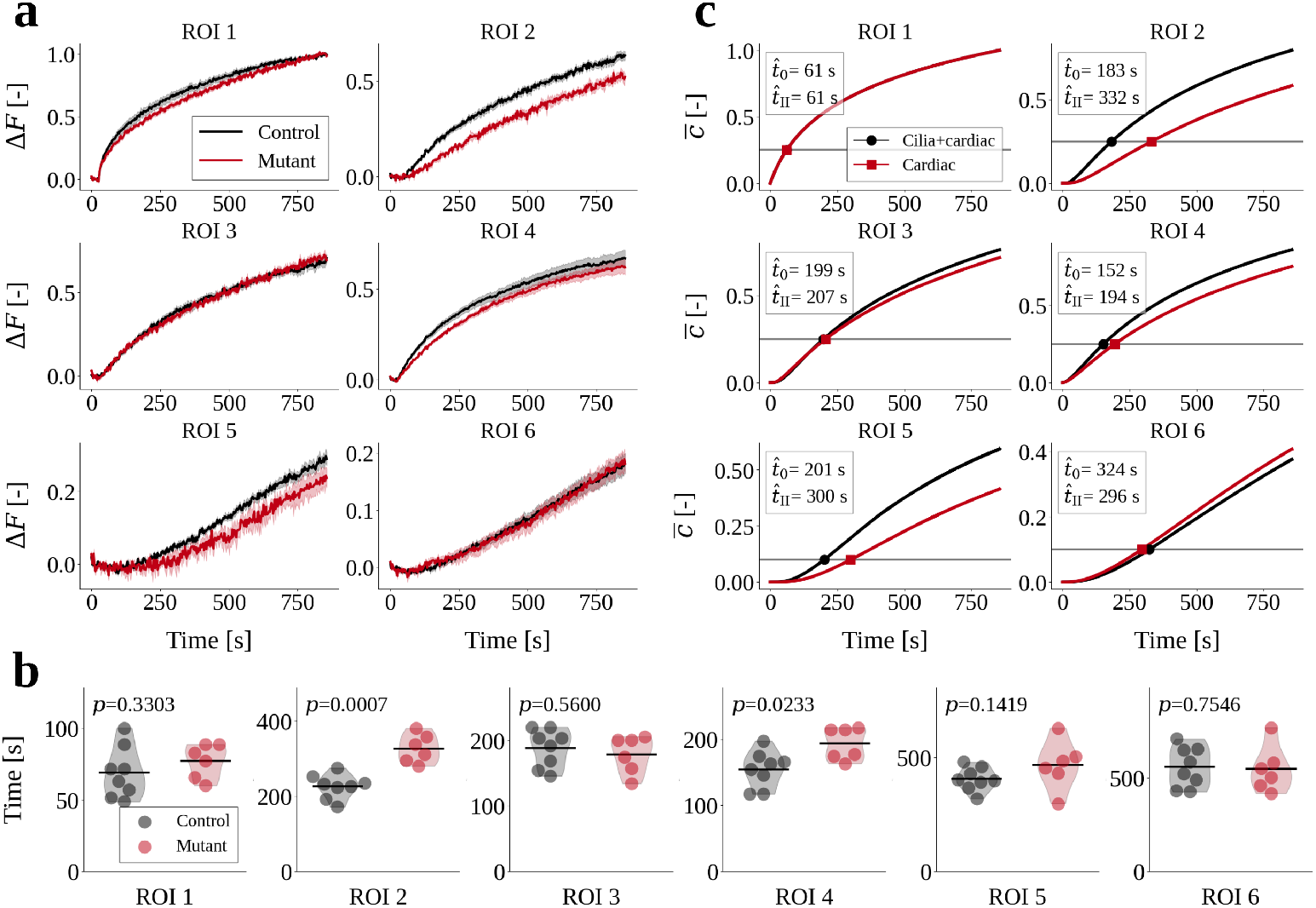
**a**. Relative change in fluorescence intensity Δ*F* (*t*) in each region of interest (ROI) for control (black lines) and *smh* mutant (red lines) zebrafish. Bold lines are mean values of the cohorts and shaded regions cover one standard deviation. **b**. Times when Δ*F* (*t*) first exceeds a threshold value, equal to 0.25 (ROIs 2–4) or 0.10 (ROIs 5, 6), for controls (gray dots) and *smh* mutants (red dots). The thick black lines denote mean values. The *p*-values were calculated using a Mann-Whitney U test with 95% confidence level, under the null hypothesis that the two distributions of control and mutant data are from the same population. **c**. Mean concentration 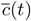 in each ROI, simulated using the baseline model (black lines) and the cardiac-induced/no-cilia model (red lines). The horizontal gray lines mark the threshold value *ĉ*, and the time 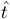 when the threshold is first exceeded is reported in the plot legends. Note that in panels **a** and **c** the vertical axes have different limits in each plot.

In the computational model, we represent the *smh* mutants by considering transport of Dendra2 with CSF flow driven by cardiac pressure pulsations only, representing the absence of motile cilia. Comparing the in silico mutants with transport predictions in the presence of both cardiac pulsations and motile cilia, we observe noticeably delayed dynamics in ROIs 2, 4 and 5, but little change for ROIs 3 and 6 (Figure 4c).

Compared to experimental data, the control and mutant scenarios differed more for the simulations, with the times-to-threshold in ROI 2 and 4 being 81% and 28% higher in the absence of cilia-mediated flow. Moreover, the predicted mean concentrations in ROI 5 deviate between the two scenarios, while in the experiments the difference in fluorescence intensities is not significant. Altogether, our findings identify that, for smaller proteins such as Dendra2, diffusion plays an important role for solute movement to regions upstream in the cilia-mediated flow fields or to distant ventricles, while the cilia affect the downstream mean concentrations.

### 2.5 Regional loss of cilia motility delays transport

We previously characterized three distinct ciliated cell lineages in the zebrafish larval brain.^[14, 18]^ To identify the effects of these lineages on CSF flow and transport, we investigate region-specific cilia paralysis with the computational model. Three cilia-covered regions were paralyzed separately: the anterior ventricle, and the dorsal and ventral parts of the middle ventricle (Figure 5a). To simulate regional cilia paralysis, we remove the tangential traction exerted by the cilia in the respective region.

**Figure 5:**
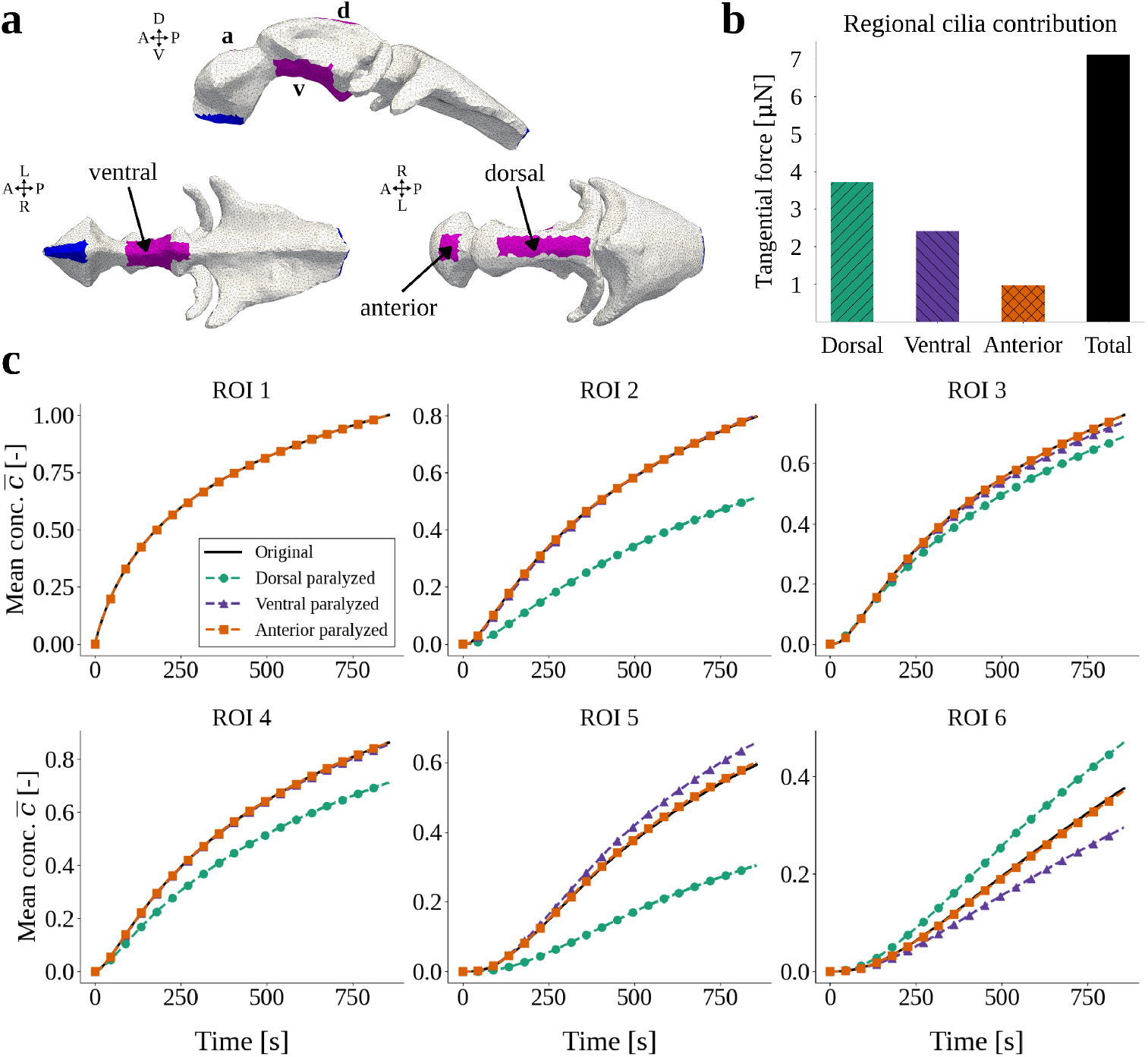
**a**. The three separate cilia regions for which cilia paralysis is simulated by removing the cilia tangential traction boundary condition in the respective region. **b**. The total force applied to the boundary with the cilia traction in each of the three separate ciliated regions, and the total force (the sum of the three regions), calculated with Equation (6). **c**. Mean concentration 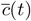 for the six regions of interest (ROIs) in the three paralysis scenarios. The legend refers to the region where cilia are removed.

Removing cilia in the dorsal part of the middle ventricle has a strong effect on the CSF velocities: the maximum velocity magnitude drops from 27.9 µm/s to 7.9 µm/s. In vivo, these dorsal cilia generate the strongest flow at this developmental stage of the fish (Figure 1c).^[14]^ In contrast, only removing cilia in the anterior ventricle or the ventral region of the middle ventricle has minimal impact, and results in maximum speeds of 27.9 µm/s and 27.4 µm/s, respectively. We remark that out of the total tangential forces, ventral cilia in the middle ventricle contribute by 34%, while the dorsal cilia exert 52% of the total tangential force (Figure 5b). Thus, the CSF velocities are affected not only by the magnitude of the applied forces, but result from a nontrivial interplay between force directionality and location, and ventricular morphology.

Turning to transport of Dendra2, removal of the dorsal cilia again has the most pronounced effect. All mean concentration profiles (except ROI 1, where we impose the photoconversion curve) change noticeably (Figure 5c). Compared to the baseline model results, we observe less solute distribution in the middle and anterior ventricles (ROIs 2–5), and more rapid transport to the posterior ventricle (ROI 6). On the other hand, removal of anterior cilia does not alter the mean concentrations (Figure 5c). Finally, when removing ventral cilia in the middle ventricle more solute spreads towards the anterior ventricle (ROI 5), and less solute spreads towards the posterior ventricle (ROI 6).

### 2.6 Modification of ventricular geometry impacts solute distribution

Variations in ventricular morphology impact CSF flow patterns, and we previously identified large interindividual variation in ventricular size and CSF velocities between larva at given developmental stages and throughout development.^[14, 18, 58]^ To assess how these variations influence solute distribution, we consider four alternative ventricular geometries: (i) constriction of the anterior-middle interventricular duct (66 % reduction in cross-sectional area); (ii) constriction of the middle-posterior duct (33 % reduction); (iii) middle ventricle shrunk in the lateral direction (43 % reduction); and (iv) all three combined (Figure 6a, b). In these four modified geometries, the total volume is reduced by 3.9%, 4.1%, 1.4% and 9.4%, respectively, compared to the original geometry. As a result of the geometry modifications, the forces applied with the cilia tangential traction differ across the scenarios (Figure 6c).

**Figure 6:**
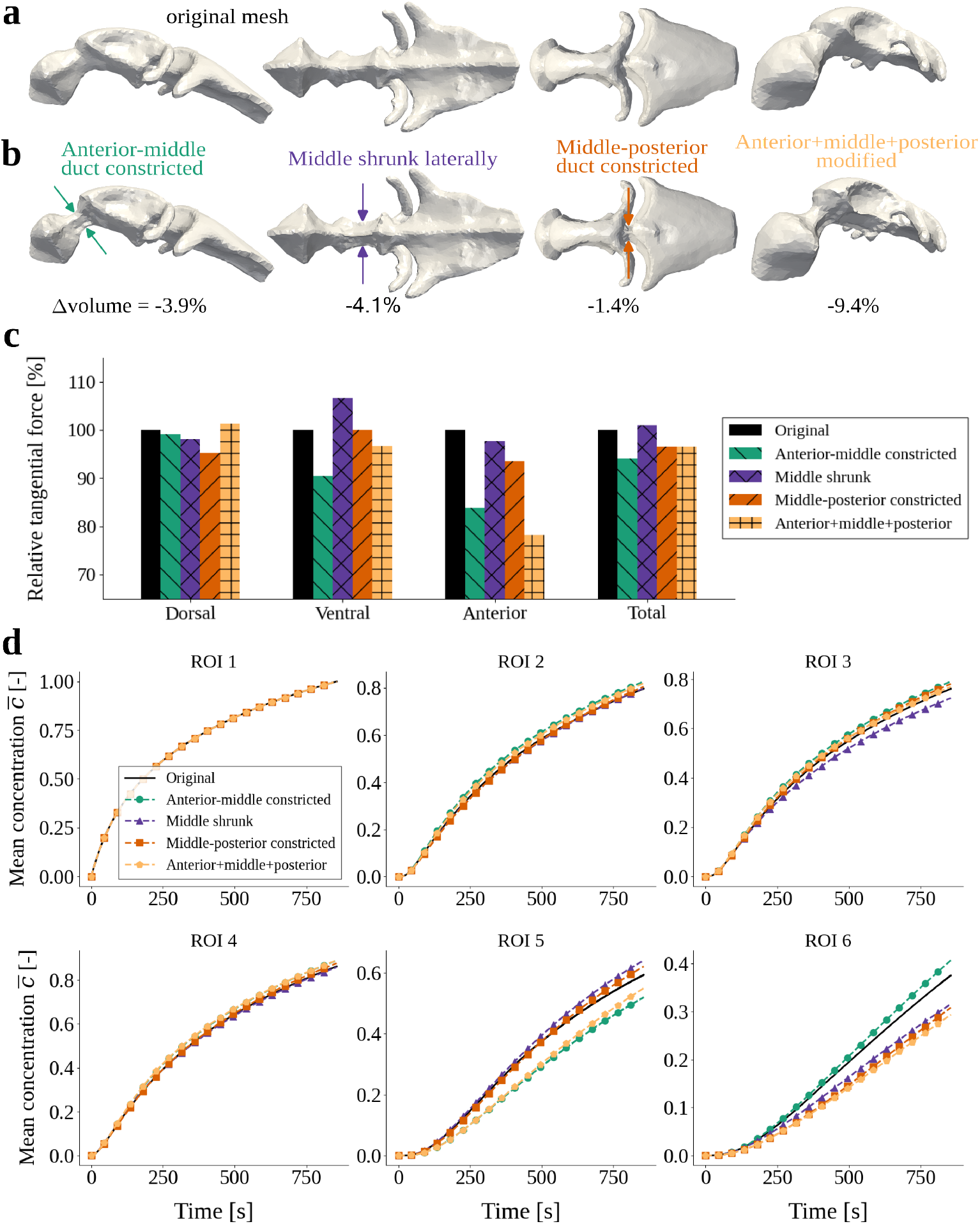
**a**. The original computational mesh of the ventricular geometry. **b**. The four modified versions of the computational geometry. Compared to the original mesh, the volumes of the deformed meshes are reduced by 3.9%, 4.1%, 1.4% and 9.4%. **c**. Mean concentration *c*(*t*) simulated with the original and modified geometries in the six regions of interest (ROIs). **d**. A comparison of the tangential cilia force in the three cilia regions and the total force applied, calculated with Equation (6). In each region, the force *F*_original_ applied on the original mesh is the baseline of 100%, to which the forces *F*_*i*_ of the other mesh versions are compared, where *i* denotes the modification scenario. The bars for the four alternative scenarios display *F*_*i*_*/F*_original_ × 100%.

Constricting the anterior-middle duct reduces solute transport to the anterior ventricle (ROI 5), and the mean concentrations in ROIs 2–4 and 6 slightly increase, compared to the original geometry (Figure 6d). Constricting the middle-posterior duct mainly delays the transport into the posterior ventricle (ROI 6), while slightly accelerating transport towards the anterior ventricle (ROI 5). Thus for both scenarios, solute transport is restricted through the respective constricted interventricular duct. For the geometry with a shrunk middle ventricle, we observe faster transport to the anterior ventricle (ROI 5) and slower transport to the lower middle and posterior ventricles (ROI 3, ROI 6). Interestingly, this reduction in the transport to the posterior ventricle (ROI 6) happens in spite of a 6.6% increase in the ventral cilia forces, which are the cilia that propel CSF towards the posterior ventricle.

When combining all three modifications, we observe higher retention of solute in the middle ventricle (ROIs 2–4), while the mean concentrations are reduced in both the anterior and the posterior ventricles (ROIs 5 and 6). To summarize, these results show that the ventricular geometry and the cross-sectional area of the ventricular ducts clearly impact solute transport, with notable differences in transport resulting from modest geometry modifications. These findings may also explain the larger differences in the cilia-deficient computational model results compared to the differences in experimental data in Figure 4, as the computational geometry is not an exact representation of the individual zebrafish ventricles.

## 3 Discussion

In this study, we have presented a computational model of CSF flow and solute transport within brain ventricles. We used the larval zebrafish brain ventricles since their geometry and flow mechanisms are well described, thereby allowing for validation of the simulation results against experimental measurements. Our computational framework can be adapted to other ventricular systems, once their flow parameters are adequately resolved. Here, we assumed that intraventricular CSF flow was governed by the Stokes equations. By imposing tangential traction and normal pressure acting on the CSF, we were able to reproduce flow features observed experimentally. In particular, vortex-structured flow mediated by the cilia and pulsatile flow mediated by the cardiac cycle emerged as key flow characteristics.

Using the simulated CSF flow, we modeled solute transport via an advection-diffusion equation. The resulting evolution and distribution of solutes compared well with data from in vivo experiments in zebrafish and is advection-dominated on the global scale for a physiologically-relevant range of solutes. By studying the distribution patterns in the absence of cilia motion, both via zebrafish mutants and computationally, we find that diffusion more strongly affects solute spread to distant parts of the ventricular system (upstream). Regional loss of cilia delayed solute transport with the most pronounced effect stemming from dorsal cilia paralysis. Moreover, the ventricular morphology also affects solute distribution, with the duct between the anterior and middle ventricles as a key passage. Altogether, we present not only new findings on the roles of advection and diffusion in brain ventricular transport, but also provide a computational framework to study CSF flow and transport in ventricular geometries.

### 3.1 The roles of advection and diffusion in brain ventricular solute transport

We quantified brain ventricular solute transport based on changes in fluorescence intensity Δ*F* (experiments) and mean concentration 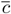 (simulations) over time. Our results show that the solute distribution depends on a balance between advection and diffusion. While advection is the strongest transport mechanism on a global scale for the three solutes we studied (global Péclet numbers *>* 1), diffusion is balancing out advection for shorter distances and in regions without motile cilia activity. For instance, for the larger extracellular vesicles, no solute reached the distant posterior ventricle (a distance of *L* = 200 µm) within the simulation time (around 860 s). Advective transport by the mean velocity 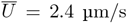 over the same distance would occur on a timescale of 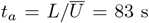, indicating that transport to the posterior ventricle is not driven by advection at this developmental stage. The estimated timescale of diffusion for the extracellular vesicles (*t*_*d*_ = *L*^2^*/D*_1_) over the same distance is 24500 s, but as low as 696 s and 348 s for the smaller proteins STM-GFP and Dendra2, respectively. These estimates support our observations that STM-GFP and Dendra2 spread to the posterior ventricle (by diffusion) while the extracellular vesicles do not. Larger molecules transported primarily by advection are thus more easily compartmentalized within the ventricles as a result of the compartmentalized CSF flow.

### 3.2 Effects of cilia motility on transport

Removing motile cilia delayed transport within the middle ventricle and towards the anterior ventricle, while the transport to the posterior ventricle was unaffected. Photoconversion experiments with cilia mutant zebrafish larva (*smh*) aligned with these observations. These simulations and experiments further indicate that transport from the middle to the posterior ventricle is purely diffusive at this developmental stage, while transport within the middle ventricle depends on cilia motility. Furthermore, our experimental results provide quantification of the impact of cilia motility on brain ventricular transport. Interestingly, Yoshida *et al*. observed a drastic reduction in CSF mixing in specific regions of the human brain ventricles when simulating reduced cilia motility,^[52]^ an observation that compares well with our findings.

### 3.3 Model validation

Our computational predictions and in vivo experimental measurements agree to a large extent. Examining any discrepancies in more detail indicates that the simulated transport dynamics are more rapid and results in higher total amounts of solute. Differences between experiments and simulations may relate to ventricular size, as we showed that modifying the ventricular geometry impacts solute distribution, and the ventricular size varies substantially in the control population at 2 dpf. In addition, differences may result from the definition of the photoconversion site, since the volume of this region determines the total amount of solute and thereby the rate of transport. Another source of discrepancy may relate to the permeability of the ventricular wall, which we considered as impermeable in our simulations. The neuroepithelium lining the walls of brain ventricles in 24 hour old zebrafish is permeable to dyes smaller than 70 kDa.^[68]^ If this is also the case at 2 dpf, Dendra2 (which are 26 kDa in size^[69]^) would leak out of the brain ventricles over the relevant time spans and render the dynamics slower. It may also be the case that our model predicts faster transport because the Dendra2 diffusion coefficient we used (based on an experimental value^[70]^) is too high. This could be because embryonic CSF has higher viscosity than the solution considered in the measurements, or because the presence of proteins, nutrients and other solutes obstructs diffusive transport.^[71–73]^ Finally, an important difference is the three-dimensional nature of the computational model, while the experimental images only show 2D slices.

We modeled motile cilia as a tangential traction at the ventricular walls while allowing the CSF to slip, motivated by experimental observations of flow fields in zebrafish larval brain ventricles in which the fluid velocity increases towards the wall (Figure 1c).^[14]^ The ciliated boundaries in our model are regionally homogeneous, whereas in reality cilia populate the ventricle walls heterogeneously.^[14]^ Heterogeneity in cilia location, beating direction and beating frequency enhances particle clearance in airway cilia arrays in mice.^[54]^ Beating directionality can affect transport, and specific coordination of the ciliary beating is important for flow and signaling processes in the brain ventricles,^[14, 24, 33, 55, 61, 62]^ for physiological mucous transport in airways,^[26, 74–76]^ and in the nodal flow that establishes left-right symmetry in mammals.^[42, 63, 64]^ Incorporating spatial heterogeneity in our model would therefore presumably affect results, and could be interesting to pursue in further work.

The maximum stress of the cilia boundary condition was |***τ***| _max_ = 2.5 mPa = 2.5 × 10^−3^ N*/*m^2^. In a model of CSF flow in human brain ventricles, Siyahhan *et al*. modeled ciliary motion with a force increasing linearly from the ventricular wall to a maximum force density of *f*_max_ = 526 N*/*m^3^ at a distance 15 µm (the cilium length) away.^[32]^ Their simulation results suggest that cilia only contribute to near-wall flow dynamics, which contrasts our results in which the cilia induce large-scale flow. This is a consequence of the scale of the cilia and the difference in ventricular size between humans and zebrafish. In a cilia array of thickness *δ* = 15 µm, the maximum stress in our model would correspond to a maximum force density of

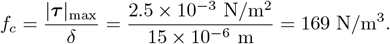

With *δ* = 5 µm (the typical cilium length in zebrafish brain ventricles^[14, 30, 53]^) our cilia stress corresponds to a maximum force density of *f*_*c*_ = 507 N*/*m^3^, similar to the value Siyahhan *et al*. used.^[32]^ For further comparison, Thouvenin *et al*. modeled two-dimensional cilia-mediated flow in the central canal of zebrafish embryos using a force density of 4000 N*/*m^2^.^[29]^ Averaging this value over the reported central canal diameter 8.9 µm yields a maximum force of 449 N*/*m^3^. In light of the previous finding that motile cilia align by response to shear stress magnitudes of flow,^[24]^ it is interesting to note the above similarity in cilia force estimates, despite the variety in application.

### 3.4 Numerical considerations

When simulating solute transport, we handled the advective transport through the anterior/posterior boundaries (where cardiac pressure pulsations were imposed) via an approximate periodic boundary condition that conserved the total solute concentration. A possible improvement would be to consider a 3D-0D coupling of the in- and outflow using ordinary differential equations, similar to applications in cardiac modeling.^[77, 78]^ Alternatively, the beating cardiac motion could be modeled by ventricular wall deformation to avoid advective boundary fluxes.^[13, 79–81]^ We also remark that the Brezzi-Douglas-Marini and discontinuous Galerkin scheme BDM_1_-DG_0_ used to discretize the Stokes equations is computationally more expensive than more common discretizations such as Taylor-Hood (*P*_2_ − *P*_1_ elements)^[82]^ or lower-order enriched elements (e.g. MINI).^[83]^ However, we prefer the BDM-DG scheme because of its mass conservation properties: the numerical approximation of the CSF velocity ***u***_*h*_ satisfies the continuity equation (Equation (1b)) point-wise, an important property for the stability of the advection-diffusion equation, especially in advection-dominated cases.^[84–86]^ Additionally, the BDM-DG scheme lets us impose the impermeability condition ***u*** · ***n*** = 0 directly^[87]^ (see Appendix A.1 for more details).

### 3.5 Outlook

Altogether, we have shown that the flow generated by ciliary motion in the zebrafish larval brain ventricles is of the scale of the geometry.^[14]^ In contrast, one can expect that in large brains such as the human brain, ciliary beating contributes only to flow proximal to the ventricular walls.^[32]^ Nevertheless, the cilia may still play an important role in regulating the amount of substances, especially extracellular vesicles, close to the ventricular walls, similar to coral reefs where cilia are primary regulators of oxygen amounts at the coral surface.^[20]^ Notably, the discovery of a cilia-driven flow network in rodent brains strongly supports such selective brain ventricular transport.^[33]^ Our results also revealed that solute concentrations are affected by ventricular geometry changes. In fact, a 9% difference in volume had a noticeable impact on solute transport, suggesting that pathological ventricular volume changes, such as the enlargement observed in hydrocephalus, could affect CSF distribution and composition. Because CSF plays an important role in the regulation of solutes within the ventricular system,^[88]^ understanding more of the interplay between transport dynamics and ventricular morphology, as well as the impact of reduced mixing resulting from cilia-deficiency, are important considerations for future work.

## 4 Methods

### 4.1 Zebrafish maintenance, genotyping and strains in photoconversion experiments

The animal facilities for zebrafish (*danio rerio*) are approved by the Norwegian Food Safety Authority (NFSA, Mattilsynet). The zebrafish were maintained in accordance with the guidelines set by the NFSA and the European Communities Council Directive.

The larval and adult zebrafish were raised under standard husbandry conditions at 28.5°C in a Techniplast Zebtech Multilinking system. The fish tanks were kept at constant pH 7.0 and 685 µS with a 14/10 hr light/dark cycle. From fertilization to 3 days post-fertilization (dpf), larvae were maintained in egg water (1.2 g marine salt and 0.1% methylene blue in 20 L reverse osmosis (RO) water) and subsequently transferred to artificial fish water (AFW) (1.2 g marine salt in 20 L RO water). The zebrafish lines used in the experiments were *ccdc103(dnaaf19)*^*tn222a*^ (*schmalhans (smh)*)^[89]^ and *Tg(gfap:Gal4FF)*^*nw7Tg*^,^[90]^ *Tg(5xUAS:Signal-Dendra2)*^*nw21Tg*^. Animals were in the pigmentless *nacre*^*b692*^ (*mitfa*^*-/-*^)^[91]^ background.

Wholemount in vivo live imaging and photoconversion experiments were performed with zebrafish larvae at the 2 dpf stage obtained from inbreeding of heterozygous *smh*^*+/-*^;*Tg(gfap:Gal4FF);Tg(5xUAS:Signal-Dendra2*) adult animals, as described by D’Gama *et al*.^[18]^ Controls consisted of either wild-type (*smh*^+/+^) or *smh*^+/-^ heterozygous zebrafish from the same breeding. The mutants were identified based on their curved body.^[89]^

For genotyping, genomic DNA (gDNA) was isolated from clipped fins of anesthetized adult fish using 100 µL of PCR lysis buffer (containing 1M tris pH 7–9, 0.5 M EDTA, tritonX-100, and Proteinase K 0.1 mg/ml) overnight at 50°C. To stop the lysis reaction, the samples were heated to 95°C for 10 minutes and then centrifuged at 13000 rpm for 2 minutes. The supernatant containing gDNA was utilized for KASP assays-based analysis. The gDNAs were diluted (1:2) with water. The KASP assay was run following the guidelines of the manufacturer (LGC Biosearch Technologies−) with 3 µL gDNA, 5 µL of master mix, 0.14 µL of assay mix and 1.86 µL of RO water per well on a 96-well plate.

### 4.2 Generation of transgenic line to study CSF and transport dynamics

To generate a transgenic line expressing the photoconvertible protein Dendra2 in the brain ventricles, we fused the open reading frame (ORF) of the neuropeptide y (npy) signal peptide (*npy* : ENSDARG00000036222, Q1LW93) to the N-terminal of zebrafish codon optimized Dendra2 DNA sequence. The DNA sequence was synthesized with EcoR1 enzyme sites at the 5’ and 3’ ends (GenScript Biotech) and inserted into the pT2MUASMCS vector^[92]^ through restriction enzyme cloning (GenScript Biotech) to generate the 5xuas:Signal-Dendra2 plasmid DNA.

A volume of 2 nl of a mixture of the plasmid DNA (60 pg) and tol2 mRNA (10 pg) was microinjected into one-cell stage embryos to generate the transgenic line, as described in Jeong *et al*.^[58]^ The injected embryos were raised to adulthood (F0). Germline-transmitted founder zebrafish were identified by breeding with multiple Gal4 transgenic lines. Stable F1 embryos expressing the Gal4-driven Dendra2 signals were screened and raised to adult zebrafish.

### 4.3 Wholemount zebrafish in vivo time-lapse imaging and photoconversion of Dendra2

Two dpf larval zebrafish were anesthetized in 0.013% MS222 (Ethyl 3-aminobenzoate methanesulfonate, Sigma) in AFW and mounted laterally in 1.5% low melting agarose in a Flurodish (VWR, FD35PDL-100), to obtain a lateral view of the brain ventricles. After the fish were positioned in the agarose, the dish stood for 5 min to allow the agarose to solidify. After solidification, AFW containing 0.013% MS222 was added, and then the fish were transferred to the confocal microscope (LSM880 Examiner, Zeiss) and imaged using a 20x water-immersion objective (Zeiss, NA 1.0, Plan-Apochromat) at room temperature.

The time-lapse images were acquired in a single plane, covering the telencephalic, diencephalic and rhombencephalic ventricles simultaneously, with a frequency of 0.37–0.38 Hz (2.60–2.68 sec / frame). The size of the images was 1024 × 400 pixels. In total, 300 images were acquired per fish. The first 10 images were scanned without photoconversion to obtain a baseline value of fluorescence intensity, and the next 290 images were scanned while performing photoconversion.

For photoconversion, “Bleaching” and “Region” in the ZEN software were used. A 405 nm laser was focused into a circular area (diameter: 16 µm, scan 0.77 µsec/pixel) of the dorsoposterior diencephalic ventricle with 100% laser power. The laser was illuminated in the area repeatedly between each scanning. After imaging, the fish health was checked and only data from healthy fish were analyzed. Data were collected and analyzed from three separate experiments, with a total of 8 controls and 6 mutants.

### 4.4 Photoconversion data analysis

The image data analysis was performed in MATLAB and the codes are openly available.^[93]^ The photoconverted channel was aligned to correct for drift in *x, y* directions using previously developed codes.^[28, 30]^ Only stable recordings were further analyzed. To increase the signal-to-noise ratio, the images were downsampled by a block average using a resampling factor of 6. The change in fluorescence intensity Δ*F* = (*F* − *F*_0_)*/F*_0_ was calculated for each pixel and each time point. Here, the value *F*_0_ is the average fluorescence intensity for the baseline (the first 10 time-lapse images) acquired before photoconversion.

To identify the time needed to pass a certain threshold value, Δ*F* values were first smoothened. The first timepoint when values surpassed the threshold was reported per pixel (in the downsampled images). A mask for the image was generated using the green (not photoconverted) channel and calculated on the block-averaged data based on the intensity being higher than 1.5x median intensity of all the frames.

To obtain Δ*F* values for each region of interest (ROI), ROIs were first drawn on the aligned and block-averaged time series (Figure 2). Six ROIs were drawn manually: ROI 1 around the location of photoconversion, and ROIs 2–6 in different regions of the ventricular system. The pixel values of the fluorescence intensity within one ROI were averaged for each time point, and the relative change Δ*F* calculated with this averaged value. To normalize the data with respect to photoconversion efficiency, the Δ*F* values for all ROIs were divided by the Δ*F* values obtained for ROI 1 at the end of the experiment, so that Δ*F* values for the photoconverted site ROI 1 ranged from zero to one. Finally, we averaged the Δ*F* data from controls and from mutants.

### 4.5 Intraventricular injection of microbeads and particle tracking

We carried out particle tracking in a larval zebrafish injected with fluorescent beads, as described in previous work.^[14]^ Anesthetized 2 dpf zebrafish were injected with 1 nl of a mixture containing 0.1% w/v fluorescent beads (SPHERO Fluorescent Yellow Particles 1% w/v, F = 0.16 mm) diluted in 7.5 mg/ml 70 kDa rhodamine B isothiocyanate-dextran (RITCdextran; Sigma-Aldrich, R9379) dissolved in artificial CSF. Artificial CSF composition was as follows: 124 mM NaCl, 22 mM D-(+)-Glucose, 2.0 mM KCl, 1.6 mM MgSO_4_ · 7 H_2_O, 1.3 mM KH_2_PO_4_, 24 mM NaHCO_3_, 2.0 mM CaCl_2_ · 2 H_2_O. The needles used for the injections were pulled with a Sutter Instrument Co. Model P-2000 from thin-walled glass capillaries (1.00 mm; VWR) and cut open with a forceps. A volume of 1 nl of the solution was injected with a pressure injector (Eppendorf Femtojet 4i) in the rostral rhombencephalic ventricle. The pressure and time were calibrated for each needle using a 0.01 mm calibration slide.

Following injection, the zebrafish was directly placed under the confocal microscope and 1200 images were acquired at a frequency of 13.1 Hz. A single optical section was obtained. Particle tracking was performed using TrackMate^[94]^ in Fiji/ImageJ^[95]^ and plotted using MATLAB, as described by Jeong *et al*.^[40]^ The parameters for TrackMate were as follows. The DoG detector was used with the settings: threshold: 2.0, with median filtering; radius: 2.0, with subpixel localization. Next, the Simple LAP tracker was used with settings: max frame gap: 2; alternative linking cost factor: 1.05; linking max distance: 5.0; gap closing max distance: 2.0; splitting max distance: 15.0; allow gap closing: true; allow track splitting: false; allow track merging: false; merging max distance: 15.0; cutoff percentile: 0.9. Only the particle tracks with at least 30 data points, while simultaneously covering a distance of at least 8.8 µm, are shown (Figure 1b, c).

### 4.6 Image-based computational geometries of zebrafish brain ventricles

To model CSF flow and transport in the zebrafish brain ventricles, we generated a 3D surface representation of the ventricular walls using confocal imaging data of a zebrafish embryo injected intraventricularly with a 70 kDa dye at 2 dpf.^[14]^ At 2 dpf, the ventricular system consists mainly of 3 cavities: the telencephalic (anterior), the di-/mesencephalic (middle) and the rhombencephalic (posterior) ventricles, which are connected by interventricular ducts. From the surface geometry, we constructed a volumetric mesh of the ventricles by initially using fTetWild^[96]^ and subsequently local refinement functionality in DOLFINx.^[97]^ The geometry spans approximately 600 microns along the rostrocaudal axis and around 300 microns in the lateral direction. At most around 100 microns separates the ventral and dorsal ventricular walls. The standard mesh consists of 141 512 tetrahedral cells with a maximal (minimal) edge length of 14.9 µm (3.1 µm) (Figure 1d). Parts of the ventricular surfaces were marked to be lined with motile cilia (Figure 1d, magenta markers).^[14]^

The morphology and geometry of the brain ventricles vary across zebrafish individuals, and under physiological and pathological conditions.^[14]^ Motivated by an interest in how ventricular geometry impacts solute distribution, we also consider four variations in the geometry (cross-sectional area reduction in parentheses): by shrinking the fore-mid brain connection (66 %), the middle ventricle (33 %), the mid-hind brain connection (43 %), and all three simultaneously. We used Blender^[98]^ to modify the geometries (Figure 6a, b).

### 4.7 Computational CSF dynamics in the brain ventricles

The beating motion of the cilia generates CSF flow with persistent rotational structures^[14]^ within the brain ventricles (Figure 1a–c). In addition, cardiac pulsations induce pulsatile CSF flow. We model this flow of CSF by the time-dependent, incompressible Stokes equations which read as follows. Find the CSF velocity ***u*** = ***u***(***x***, *t*) = (*u*_*x*_(***x***, *t*), *u*_*y*_(***x***, *t*), *u*_*z*_(***x***, *t*)) and the CSF pressure *p* = *p*(***x***, *t*) for ***x* ∈** Ω and time *t* ∈ (0, *T*], such that

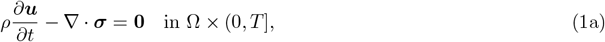

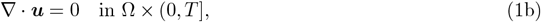

where the stress tensor is defined as ***σ***(***u***, *p*) = 2*µ****ε***(***u***) − *p***I** and 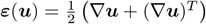 is the strain-rate tensor. Bold-face characters denote vectors or tensors, and **I** is the identity tensor in three dimensions. Furthermore, *ρ* = 1000 kg*/*m^3^ is the CSF density and *µ* = 0.7 mPa · s is the dynamic viscosity of the CSF.^[71]^ We introduce the tangential traction 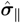 as the tangential component of the traction 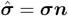:

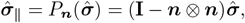

where ***n*** is the outer unit normal of the boundary surface Γ = *∂*Ω, and *P*_***n***_(***r***) is the tangential projection of a vector ***r*** onto Γ. We consider three types of boundary conditions: (i) *tangential traction* (no flow normal to the boundary ***u***·***n*** = 0 with a stress ***τ*** applied tangentially on the boundary 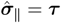), (ii) *normal pressure* (a pressure 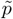 applied in the normal direction: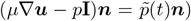, and (iii) *free slip* (no flow normal to the boundary ***u· n*** = 0 and no tangential traction 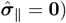. We assume that the system starts at rest: ***u***(***x***, *t* = 0) = **0**.

To study the impact of the motile cilia and the cardiac pulsations on the CSF flow, we consider the following computational flow scenarios, each predicting a CSF flow velocity ***u*** and pressure *p*. In all of the scenarios, Γ_s_ denotes a free-slip boundary.

- A *baseline* flow model representing a wild-type zebrafish including contributions both from the cilia forces and the cardiac pulsations. In this scenario, we let Γ = Γ_s_ ∪ Γ_c_ ∪ Γ_p_ and apply the tangential traction ***τ*** on the cilia boundary Γ_c_ and prescribe the normal pressure 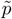 on the anterior/posterior boundary Γ_p_.
- A *cilia-driven/no-cardiac* flow model including contributions from the cilia forces but no cardiac pulsations. Here, Γ = Γ_s_ ∪ Γ_c_, and we apply a tangential traction ***τ*** on the cilia boundary Γ_c_. In the absence of cardiac-induced flow, the anterior/posterior boundary Γ_p_ = ∅.
- A *cardiac-induced/no-cilia* flow model including contributions from the cardiac pulsations but no cilia forces. In this scenario, we disregard the cilia and consider Γ = Γ_s_ ∪ Γ_p_, prescribing the normal pressure 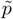 on the anterior/posterior boundary Γ_p_.

The tangential traction and normal pressure are described further below.

#### 4.7.1 Cilia-driven flow

On the ciliated regions of the ventricle walls Γ_c_ (Figure 1d, magenta markers), we impose a constant-in-time tangential traction 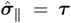 to represent the net forces of the cilia acting on the CSF. We set ***τ*** (***x, r***) = *τλ*(***x***)*P*_***n***_(***r***), where the function *λ*(***x***) and the sign of the vector ***r*** = ± (1, 0, 1) varies depending on the cilia region (see Appendix A.1). Previous experiments^[14]^ indicate flow speeds proximate to the ventricular walls of 27.4 ± 5.4 µm/s and 2.6 ± 0.6 µm/s in the dorsal and ventral regions of the middle ventricle, respectively. In the anterior ventricle, the speeds were estimated at 4.7 ± 1.3 µm/s. We calibrated ***r*** and *τ* = 0.65 mPa by comparison of the simulated maximum velocity magnitude with these flow speeds.

#### 4.7.2 Flow induced by cardiac pulsations

To generate a cardiac-induced pulsatile flow inside the brain ventricles, we set 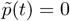 and 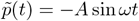 on anterior and posterior parts, respectively, of the boundary Γ_p_ (Figure 1d, blue markers). Based on a measured cardiac frequency of *f* = 2.22 Hz,^[14]^ we set the angular cardiac frequency *ω* = 2*πf* = 6.97 rad/s. The amplitude *A* = 1.5 mPa determines the magnitude of the pulsatile flow and was calibrated based on previous experiments.^[14]^ Particle tracking data were used to determine the contributions of the cardiac pulsations to the velocity magnitude, as compared to the effect of cilia motion on the flow (cf. Appendix C).

### 4.8 Computational model of solute transport after photoconversion

We simulate transport of photoconverted proteins within the brain ventricles by modeling transport of a solute with concentration *c*(***x***, *t*) for ***x*** ∈ Ω and time *t* ∈ (0, *T*] by the advection-diffusion equation

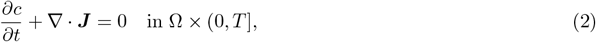

where the total concentration flux ***J*** is assumed to consist of an advective and a diffusive flux:

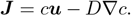

Here, ***u*** is the CSF velocity field, governed by the Stokes equations (Equation (1)), and *D* is the molecular diffusion coefficient of the solute. The ventricular walls are assumed to be impermeable (admitting *no flux* : ***J · n*** = 0), except at the anterior/posterior boundary Γ_p_. At the anterior/posterior boundary, we permit the solute concentration to be advected into and out of the domain in a manner that ensures conservation of mass (see Appendix A.3 for details).

To computationally represent protein photoconversion in a region Ω_pc ⊂_ Ω (corresponding to ROI 1), we prescribe a given time-dependent value to the solute concentration in this region. Mimicking the experimental protocol, we consider the dorsal part of the diencephalic ventricle as the subdomain Ω_pc_, and set

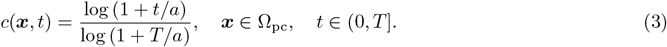

The value *a* = 65 was chosen to fit the fluorescence intensity curve observed in the physical photoconversion experiments. Because steep gradients in the concentration field may arise from imposing *c*(***x***, *t*) in Ω_pc_, the mesh is locally refined around this region.

In order to impose a smooth initial condition, we initially solve the diffusion equation

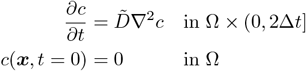

with an increased diffusion coefficient 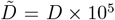 for two timesteps. The solutions are normalized by their respective maximum values, and subsequently multiplied with the value of Equation (3) at the times *t* = Δ*t* and *t* = 2Δ*t*, respectively. The resulting concentration fields are used as initial conditions for *c*(***x***, 0) and *c*(***x***, Δ*t*) for all ***x* ∈** Ω, where Δ*t* is the timestep. The two initial conditions are required because of the two-step temporal discretization scheme employed (see Section 4.12 for more details.)

### 4.9 Estimation of diffusion coefficients via the Stokes-Einstein relation

Motivated by an interest in molecules that are significant in neural development, we study the transport of molecules resembling (i) the Dendra2 Fluorescent Protein (Dendra2) used in the photoconversion, (ii) the Starmaker+Green Fluorescent Protein (STM-GFP) reported by Jeong *et al*.,^[58]^ and (iii) extracellular vesicles. To estimate their diffusion coefficients, when not available in the literature, we use the Stokes–Einstein relation

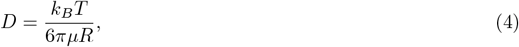

which is an expression for the diffusion coefficient of a spherical molecule suspended in a viscous fluid.^[72]^ In Equation (4), *k*_*B*_ = 1.38 × 10^−23^ J/K is the Boltzmann constant, *T* = 310 K is the absolute temperature, *µ* is the dynamic viscosity of the CSF, and *R* is the radius of the molecule. For the extracellular vesicles, we calculate *D* using Equation (4) with the relatively large *R* = 150 nm.^[99]^ For STM-GFP, we estimate the diffusion coefficient by extrapolating the experimentally observed diffusion coefficient of GFP,^[100, 101]^ assuming that if *D* scales with 1*/R* according to Equation (4), *D* would scale with the cube root of the molecule mass.^[102]^ The resulting diffusion coefficients are *D*_1_ (extracellular vesicles), *D*_2_ (STM-GFP) and *D*_3_ (Dendra2) as given by Table 1.

**Table 1:** Molecular mass and diffusion coefficients of simulated solutes. STM and GFP denote Starmaker Protein and Green Fluorescent Protein, respectively. (*) The value was extrapolated based on the value of *D* for GFP reported in the literature,^[100, 101]^ assuming that *D* scales with the cube root of the molecule mass. (†) The value was estimated using the Stokes-Einstein relation Equation (4).

| Molecule | Label | Mass [kDa] | $D$ [$\text{m}^2/\text{s}$] | Reference |
| --- | --- | --- | --- | --- |
| Extracellular vesicles (radius of 150 nm) | $D_1$ | — | $2.17 \times 10^{-12}$ | † |
| GFP (saline aqueous solution) | — | 26.9<sup>[103]</sup> | $8.70 \times 10^{-11}$ | [100, 101] |
| STM-GFP | $D_2$ | $66.2^{[104]} + 26.9^{[103]}$ | $5.75 \times 10^{-11}$ | * |
| Dendra2 Fluorescent Protein | $D_3$ | 25.6<sup>[69]</sup> | $1.15 \times 10^{-10}$ | [70] |

### 4.10 Quantities of interest

We calculate the mean solute concentrations

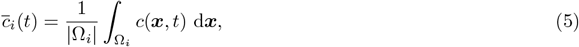

in the six regions of interest (ROIs) Ω_*i*_ (*i* = 1, …, 6) as functions of time, where |Ω_*i*_| is the volume of the domain Ω_*i*_. The dynamics of ROI 1 follows directly from Equation (3). In addition, we report the time-to-threshold as the time 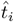 when the mean concentration 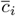 first exceeds a threshold value *ĉ*_*i*_. We use *ĉ*_*i*_ = 0.25 for ROI 1–4 and *ĉ*_*i*_ = 0.10 for ROI 5–6.

To evaluate the relative importance of advection and diffusion as transport mechanisms, we consider the Péclet number

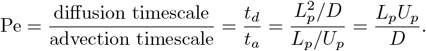

Thus, if Pe *>* 1, transport is dominated by advection, and if Pe *<* 1 transport is dominated by diffusion. Here, *L*_*p*_ and *U*_*p*_ are characteristic length and velocity scales, respectively, and *D* is the solute diffusion coefficient. In calculating a global Péclet number, we use *L*_*p*_ = 600 µm, the approximate length of the ventricular geometry along the rostrocaudal (*x*-) axis. We use a mean velocity *Ū* as the characteristic velocity *U*_*p*_. The mean velocity *U* is calculated by first averaging the velocity in space:

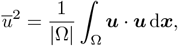

and then averaging *u* in time over one cardiac cycle to yield *Ū* = 2.4 µm/s. To assess the assumption that inertial forces are negligible compared to viscous forces, we also compute the Reynolds number

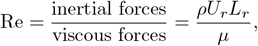

with *L*_*r*_ = 110 µm (the height of the middle ventricle) and *U*_*r*_ = 27.9 µm/s (the maximum velocity magnitude) as length and velocity scales characteristic of the flow.

When studying how cilia paralysis and ventricular morphology affect brain ventricular flow and transport, we calculate the magnitude of a tangential force *F* exerted by the cilia on a part 𝒮 ⊂ Γ_c_ of the ventricular surface as:

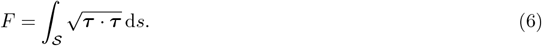

### 4.11 Numerical approximation of the Stokes equations

The Stokes equations (Equation (1)) are discretized with a finite element method in space, while in time, we employ a first-order implicit Euler discretization.^[105]^ We set the timestep size Δ*t* = 0.023 s such that 20 timesteps represent one cardiac cycle. For the spatial discretization, we use piecewise linear Brezzi-Douglas-Marini (BDM) elements for the velocity.^[87, 106]^ The BDM elements degrees of freedom are associated with integral moments of the velocity component normal to the facets of elements, thus we enforce the impermeability condition ***u · n*** = 0 strongly. We impose the boundary conditions for both the tangential traction 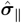 and the normal pressure 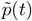 weakly. For the pressure, we use piecewise constant (zeroth order discontinuous Galerkin (DG)) elements. The resulting total number of degrees of freedom at each time step is 1 009 874. The element combination BDM_1_–DG_0_ yields a non-conforming discretization scheme for the Stokes equations.^[107]^

In particular, with BDM only the normal component of the velocity is continuous. Continuity in tangential velocities on interior facets is weakly enforced with a penalty parameter *γ* = 10.^[108]^ Since the divergence of the velocity function space is the pressure function space for this scheme, the divergence-free condition of the velocity is satisfied exactly, ensuring mass conservation.^[109]^

### 4.12 Numerical approximation of the advection-diffusion equation

The advection-diffusion equation (Equation (2)) is discretized in space with quadratic DG elements using a symmetric interior penalty DG method.^[110]^ The scheme was chosen due to its favorable mass conservation properties. At each timestep the total number of degrees of freedom is 1 415 120. In time, the equation is discretized with the second-order backward difference scheme BDF2.^[111]^ To accurately approximate the solute concentrations even in the presence of strong advection (high Péclet numbers), we use an upwind scheme for the velocity when calculating the advective flux.^[112]^ As for the flow equations, we use a timestep size of Δ*t* = 0.023 s. We simulate transport for 1900 cardiac cycles, with a final time of simulation *T* = 856 s. A DG penalty parameter *α* = 50 was chosen to ensure stability of the method.

### 4.13 Solution strategy and software implementation

Owing to the one-way coupling between the velocity field ***u*** and the concentration *c*, the governing equations for the CSF flow and the concentration can be solved sequentially. For the cilia-driven/no-cardiac flow scenario, the velocity quickly reaches steady-state since the viscous forces dominate the inertial forces. In this case, we first solve the steady-state Stokes equations (Equation (1)) for ***u*** and *p*, and then use the velocity field ***u*** as input when solving the advection-diffusion equation (Equation (2)). For the other flow scenarios, the Stokes equations are solved for one period of the pulsatile cardiac motion. The solution of this one period is then used periodically as input for the advection-diffusion equation.

The numerical methods for the Stokes equations and the advection-diffusion equation were implemented and solved with DOLFINx.^[97]^ The simulation code is openly available.^[93]^ A direct solver using MUMPS^[113]^ is used to solve the linear systems resulting from the discretized Stokes equations. For the discretized advection-diffusion equation, we use a flexible GMRES solver with block Jacobi preconditioning.^[114, 115]^ We discretize the time-dependent diffusion equation (without advection) and use the assembled linear system matrix as the preconditioner matrix for the advection-diffusion problem. Numerical experiments verified the accuracy of the model implementation (Appendix B).

## Acknowledgments

Thanks are extended to Jørgen Dokken for valuable model implementation discussions. H. H. and M. E. R. acknowledge support from the national infrastructure for computational science in Norway, Sigma2, via grant #NN8049K. M. E. R. acknowledges support from Stiftelsen Kristian Gerhard Jebsen via the K. G. Jebsen Centre for Brain Fluid Research and support from the Research Council of Norway (RCN) via grant #324239 (EMIx). N. J. Y. acknowledges support from the Research Council of Norway (grants #314189 and #353348).

## Data availability statement

The software and data used and presented in this paper, as well as the supplementary videos, are archived on Zenodo and openly available at https://doi.org/10.5281/zenodo.15194757.^[93]^

## Conflict of interest disclosure

The authors declare no conflicts of interest.

## A Supplementary theory and numerical methods

### A.1 eak formulation of the Stokes equations

We consider the time-dependent Stokes equations

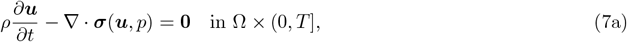

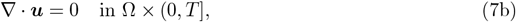

in the domain Ω with boundary Γ, where ***σ***(***u***, *p*) = 2*µ****ε***(***u***) − *p***I** is the viscous stress tensor, in which 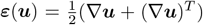 is the symmetric gradient. Furthermore, ***u*** ∈ ***V*** is the fluid velocity and *p* ∈ *Q* is the fluid pressure, and *ρ* and *µ* are the fluid density and dynamic viscosity, respectively. Unless otherwise stated, we let ***σ*** = ***σ***(***u***, *p*).

To derive weak formulations of the continuous problem defined by the Stokes equations (Equation (7)), we consider the function space^1^

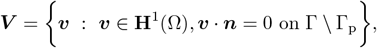

for the velocity and *Q* = *L*^2^(Ω) for the pressure. Here Γ_p_ is the anterior/posterior boundary where we impose a normal pressure boundary condition for the model versions that consider fluid motion induced by cardiac pulsations. Bold-faced characters for a space denotes a *d*–dimensional vector space, where *d* is the spatial dimension of Ω. For example, **H**^1^(Ω) = [*H*^1^(Ω)]^3^ if Ω is three-dimensional.

To derive a weak formulation of the Stokes equations (Equation (7)), we begin by multiplying Equation (7a) with a test function ***v*** ∈ ***V*** and integrating over the domain:

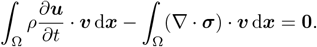

Performing integration by parts of the stress term and applying Green’s first identity, we get

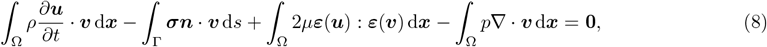

where the symbol : signifies the matrix inner product, *A* : *B* = trace(*A*^*T*^ *B*) for matrices *A, B* ∈ ℝ ^*d*×*d*^.

Further steps depend on the boundary conditions applied, which is different for the three model versions we consider in this paper. Before proceeding, let us introduce some notation used in defining the slip boundary conditions. We make use of a decomposition of any vector ***v*** on the boundary Γ into its normal and tangential components, ***v*** = ***v***_⊥_ + ***v***_∥_, where ***v***_⊥_ = (***v*** · ***n***)***n*** and ***v***_∥_ = *P*_***n***_(***v***). The tangential projection is defined as

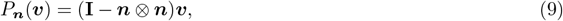

where ***n*** is the outer unit normal of the surface. Introducing 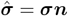 for the traction (vector), the forcing due to cilia is represented by constraining 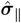 on the part of the boundary populated by cilia. In particular, we let 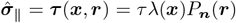, where the parameter *τ* = 0.65 mPa was used to control the magnitude of the tangential stresses, and chosen by calibrating simulated velocity fields with experimental data.^[14]^ The details of the function *λ*(***x***) follow subsequent to deriving the weak form.

We choose ***r*** = ***r***^+^ = (1, 0, 1) or ***r*** = ***r***^−^ = − (1, 0, 1) to impose forces perpendicular to the *y*-axis. The reason for this is that all of the experimental images we compare with are taken in the *xz*-plane (rostrocaudal-dorsoventral plane), rendering flow features in the *y*-direction (lateral) unobservable. In the following, we consider the way of handling the stress boundary integral for each model version successively.

#### Cilia-driven/no-cardiac model

This model version has cilia forces as the only mechanism driving flow.

We consider the Stokes equations Equation (7) with the slip boundary conditions

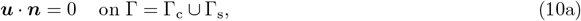

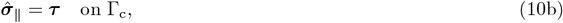

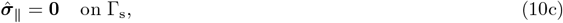

on the boundary Γ = Γ_c ∪_ Γ_s_. The boundary Γ_s_ is a free-slip boundary, where we impose the free-slip condition given by Equation (10c). The first condition (Equation (10a)) is the impermeability condition, enforcing no flow normal to the boundary, whereas Equation (10b) imposes a traction tangential to the boundary. This tangential traction is applied to represent the forces that the collective cilia motion exerts on the cerebrospinal fluid.

The boundary conditions defined by Equation (10) are set weakly through the boundary integral

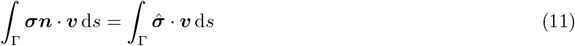

in the weak form of the Stokes equations (Equation (8)). We decompose the traction vector 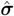 into a normal and tangential component:

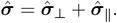

Inserting the above into Equation (11), combined with application of the tangential traction condition (Equation (10b)) and the free-slip condition (Equation (10c)), yields

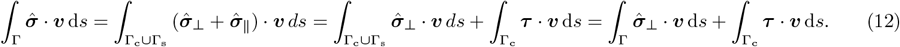

Since ***τ*** is known, the second term on the right-hand side of Equation (12) is moved to the right-hand side of the weak form. With 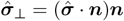, we can write the first term on the right-hand side of Equation (12) as

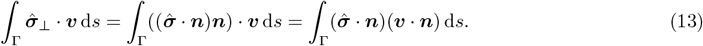

This term vanishes owing to the construction ***V***. As a result, for the cilia-driven/no-cardiac model, the weak form of the momentum equation takes the following form:

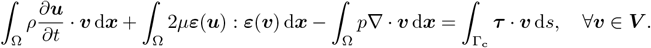

Next, multiplying the continuity equation (Equation (7b)) with a test function *q* ∈ *Q*, we require that

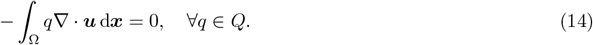

The full weak form of the Stokes equations for this flow model is then: find (***u***, *p*) ∈ ***V*** × *Q*, such that

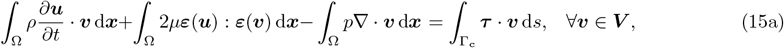

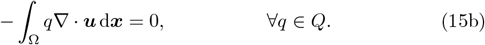

We observe that, in this case with slip boundary conditions on the entirety of the boundary Γ, for any real constant *C* the vector (***u***, *p*) = (**0**, *C*) satisfies the equations. Hence Equation (15) is a singular system with a one-dimensional nullspace of constant pressures. Consequently, assembly of the discretization of the Stokes equations (Equation (15)) into a linear equation system **Ax** = **b** yields a singular matrix **A**. To ensure existence of solutions, we orthogonalize the right-hand side vector **b** with respect to the pressure nullspace basis (the set of all constants), before solving the linear system. To obtain a unique solution, which together with existence renders the problem well-posed, information of the pressure nullspace is passed to the linear solver. Let us remark that the singularity is a consequence of the fact that the impermeability condition is assumed to hold on the *entire* domain boundary for this model version, which leaves the pressure under-determined as it is not subject to any boundary condition.

For the tangential traction ***τ*** = *τλ*(***x***)*P*_***n***_(***r***) that represents the cilia motion, we heuristically determined the constant *τ* = 0.65 mPa and the spatially varying function *λ*(***x***). The definitions of ***r*** and *λ*(***x***) for the different cilia populations are summarized in Table 2.

**Table 2:** Definitions of ***r*** and *λ*(***x***) used in the cilia tangential traction boundary condition ***τ*** = *τλ*(***x***)*P*_***n***_(***r***), where *τ* = 0.65 mPa. The *x −* coordinates *x*_0,*i*_ and *x*_*e,i*_ define the cilia domain in the dorsal (*i* = *d*) and ventral (*i* = *v*) regions of the middle ventricle.

| Cilia population | $\mathbf{r}$ | $\lambda(\mathbf{x})$ |
| --- | --- | --- |
| Anterior region, anterior ventricle | $-(1, 0, 1)$ | $\frac{2}{5}$ |
| Posterior region, anterior ventricle | $(1, 0, 1)$ | 1 |
| Dorsal region, middle ventricle | $-(1, 0, 1)$ | $\frac{11}{4} \left( \frac{x-x_{0,d}}{x_{e,d}-x_{0,d}} \right)$ |
| Ventral region, middle ventricle | $(1, 0, 1)$ | $\frac{1}{2} \left( 1 - \frac{x-x_{0,v}}{x_{e,v}-x_{0,v}} \right)$ |

#### Cardiac-induced/no-cilia model

For this model version, the only physical mechanism inducing fluid flow is the cardiac cycle. The cardiac cycle generates directional back-and-forth motion of the CSF in the zebrafish brain ventricles. We model the influence of the cardiac cycle with a normal pressure boundary condition:

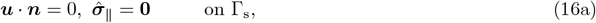

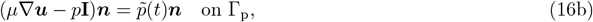

where the boundary Γ = Γ_s_ ∪ Γ_p_ consists of a free-slip boundary Γ_s_, and a part Γ_p_ where a normal traction type of boundary condition is imposed. Note, however, that the transpose velocity gradient term must be included inside the parentheses on the left-hand side of Equation (16b) for the expression to be the stress tensor ***σ***, thus the left-hand side of Equation (16b) does not constitute the normal traction 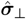. Because of this, we rather refer to this boundary condition as a normal pressure boundary condition. The boundary Γ_s_ is handled as discussed in the previous section, such that the stress boundary integral in Equation (11) vanishes over Γ_s_. Therefore, we now have

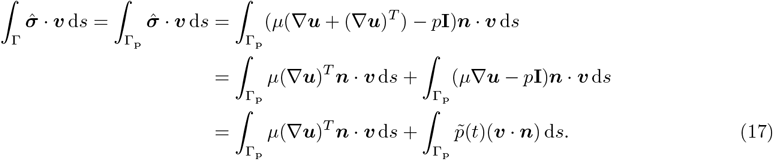

We leave the first term on the left-hand side of Equation (17) in the weak form of the momentum equation. The second term is determined by the boundary condition defined in Equation (16b), and is therefore moved to the right-hand side of the weak form. The weak form of the momentum equation is then

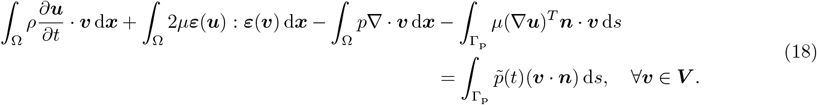

Also considering the weak form of the continuity equation, the weak formulation for the cardiac-induced/no-cilia model version then reads: find (***u***, *p*) ∈ ***V ×*** *Q* such that Equation (18) and Equation (14) hold for all (***v***, *q*) ∈ ***V*** × *Q*. To simulate the back-and-forth motion induced by the cardiac cycle observed in previous work,^[14]^ we apply sinusoidal forcing. The boundary Γ_p_ has two separate regions: one located on the anterior ventricle and one on the posterior ventricle (Figure 1d, blue markers). For the normal pressure boundary condition, we set 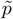 (*t*) = 0 on the anterior part and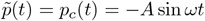 on the posterior part. The value of the amplitude *A* was based on heuristics and *ω* = 2*πf*, where *f* = 2.22 Hz is the cardiac cycle frequency.

#### Baseline model

The baseline flow model considers both cilia and cardiac beating as driving mechanisms of fluid flow. We therefore consider the boundary conditions

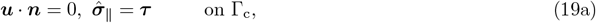

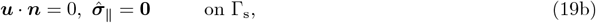

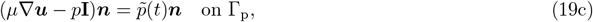

where Γ = Γ_c_ ∪ Γ_s_ ∪ Γ_p_ is the boundary of Ω. The quantities ***τ*** and 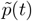 have been introduced in the previous sections covering the other flow model versions. We handle the boundary conditions Equation (19a) and Equation (19b) as introduced for the cilia-driven/no-cardiac model, and the boundary condition Equation (19c) as introduced for the cardiac-induced/no-cilia model. The abstract weak formulation for the baseline model is then: find (***u***, *p*) ∈ ***V*** × *Q* such that

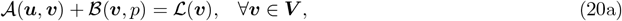

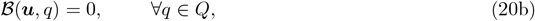

with the definitions

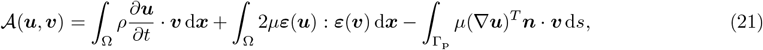

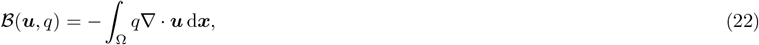

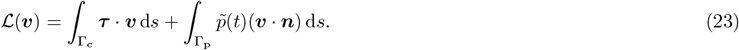

### A.2 Discretization of the Stokes equations

We outline the discretization of the weak formulation of the Stokes equations for the baseline model (both cilia traction and cardiac pressure pulsations present) given by Equations (20) to (23). A similar approach can be used to discretize the weak formulation for the other model versions. First, we consider the spatial discretization of the weak formulation. We rewrite the weak form in terms of the discretized variables, denoted with the subscript *h*, on the discrete domain Ω_*h*_:

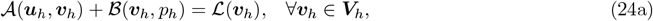

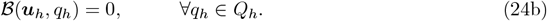

Our choice of discretization are finite elements of the Brezzi-Douglas-Marini (BDM) family^[87, 106]^ for the velocity space and discontinuous Lagrange polynomial elements for the pressure. We refer to the latter as discontinuous Galerkin (DG) elements. For all cells *K* of the computational mesh Ω_*h*_, we define

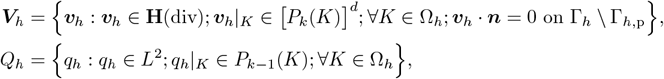

where *P*_*k*_ is the space of polynomials of degree less than or equal to *k*. The BDM family of elements have degrees of freedom associated with integral moments of vector components normal to facets of the mesh. Because ***V***_*h*_ ⊄ ***V***, this BDM-DG scheme yields a non-conforming method for the Stokes equations. The scheme is **H**(div)-conforming and exactly mass-conserving: due to the fact that ∇ · ***V***_*h*_ = *Q*_*h*_, the continuity equation (Equation (7b)) is satisfied point-wise. Moreover, control of normal components of ***u***_*h*_ conveniently lets us enforce the impermeability condition ***u***_*h*_ · ***n*** = 0 strongly.

The elements of ***V***_*h*_ only ensure by construction the continuity of normal components of ***u***_*h*_. Continuity of the tangential components of ***u***_*h*_ on interior facets of the mesh is enforced weakly by additional stabilization to the discrete weak form (Equation (24)). Specifically, we apply the stabilization outlined by Hong *et al*.^[108]^ First, we define the average and jump operators, respectively, as

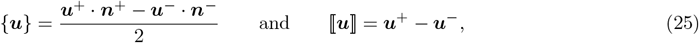

where ***u*** is a first-order tensor and ***n*** is the normal vector on a facet separating two cells, and the plus and minus signs denote variables in each of the two cells. The normal vector is chosen such that it points towards the exterior of its corresponding cell. On the boundary, the operators are defined as {***u***} = ***u***· ***n*** and ⟬***u***⟭ = ***u***, with ***n*** pointing to the exterior of the domain. Note that average operator defined in Equation (25) reduces the rank of the tensor ***u*** by one, whereas the jump operator preserves rank. This extends to tensors of higher order. We now define and add the bilinear form

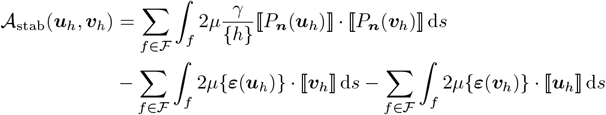

to the discrete weak formulation (Equation (24)), where F is the set of all interior facets of the computational mesh. The first term of 𝒜_stab_ penalizes jumps in the tangential components of the velocity across interior facets, where the penalty parameter *γ >* 0 must be chosen large enough to achieve stability. For appropriate convergence, the penalization term includes a mesh-dependent scaling. In this work, we choose it as the average of the cell diameter of neighboring mesh cells, with *h* defined as the maximal distance between two points in a cell.^[116]^ The other two terms of 𝒜_stab_ are added to ensure consistency and symmetry of the bilinear form, such that the problem is well-defined.

Including the stabilization outlined above, the semi-discrete weak formulation of the Stokes equations reads: find (***u***_*h*_, *p*_*h*_)∈(***V***_*h*_, *Q*_*h*_) such that

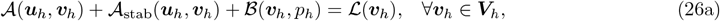

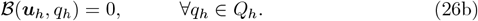

In time, we discretize the weak formulation (Equation (26)) with the implicit Euler method, approximating the time derivative as

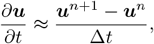

where *n* denotes the timestep index. All of the other terms in Equation (26) are evaluated at timestep *n* + 1. The method is implicit and of first order in time.^[105]^ The fully discrete Stokes equations were solved with the penalty parameter *γ* = 10.

### A.3 Weak formulation of the advection-diffusion equation

Recall the strong form of the advection-diffusion equation:

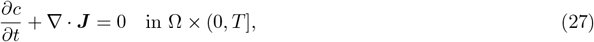

where Ω is the domain with boundary Γ, and the total solute flux

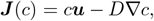

where *c*∈ *W* = *H*^1^(Ω) is the solute concentration and ***u* ∈ *V*** is the cerebrospinal fluid velocity. We consider ***J*** = ***J***(*c*), unless otherwise indicated. The weak form of Equation (27) is derived by multiplying the equation with a test function *ϕ* ∈ *W* and integrating over the domain:

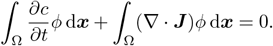

Integrating the second term by parts and applying Green’s first identity yields

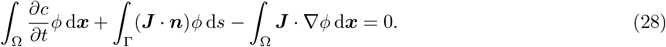

The boundary integral of the total flux is handled differently for the three different model versions, depending on the boundary conditions considered. For all model versions, we simulate photoconversion for times in Ω_pc_ ⊂ Ω by strongly enforcing

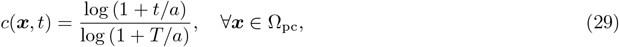

for all times *t* until simulation ends at time *T*. The parameter *a* = 65 determines the growth speed of the solute concentration and was chosen such that Equation (29) is similar to the experimental photoconversion curves.

#### Cilia-driven/no-cardiac model

We consider Ω to be a closed domain with impermeable walls, so that the whole boundary Γ constitutes a no-flux boundary Γ_nf_. By imposing ***J · n*** = 0 on Γ = Γ_nf_, the weak form defined by Equation (28) is reduced to

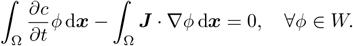

Defining the left-hand side as the bilinear form

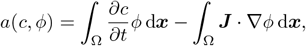

we get the following abstract weak formulation: find *c* ∈ *W* such that

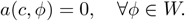

#### Cardiac-induced/no-cilia model

When using the normal pressure boundary condition (Equation (16b)) to model pulsatile flow, we have in- and outflow of cerebrospinal fluid and thus advection of *c* on the pressure boundary Γ_p_. Let Γ_p_ = Γ_in_ ∪ Γ_out_ and Γ = Γ_nf_ ∪ Γ_in_ ∪ Γ_out_, with

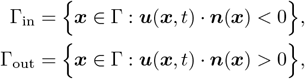

as in- and outflow boundaries, respectively, where we have emphasized that both ***u*** and ***n*** are spatially dependent variables. With ***J*** · ***n*** = 0 on the no-flux boundary Γ_nf_, we have

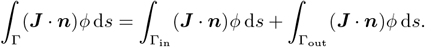

For the outflow boundary integral over Γ_out_, we impose zero diffusive flux *D* ∇*c ·* ***n*** = 0, with the advection term *c*(***u · n***) simply left in the variational form as an unknown. For the continuous problem this is problematic, since we are lacking a boundary condition to close the problem. In the discrete setting, however, this method of leaving the boundary term has been shown to yield consistent results.^[117–119]^ Intuitively, one can think of this outflow boundary condition as setting the advective flux to being the one consistent with the weak form, for which there is a unique *c* satisfying the equations, with ***u*** determined by the Stokes equations.

The flux integral over the inflow boundary Γ_in_ cannot be handled in the same way as the method used for Γ_out_, as this would not preserve coercivity of the weak form. Lacking experimental data both for the magnitude and origin of an inflow boundary flux, we deal with the inflow boundary in an explicit way that resembles an approximation of a periodic boundary condition. At a timestep *n*, when solving the linear system for the concentration *c*^*n*+1^ at the next timestep *n* + 1, we set the inflow flux 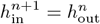, where the latter is the outflow flux

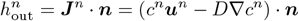

at the previous timestep. In the limit that the timestep Δ*t* → 0, we would be imposing

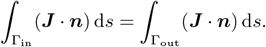

This implies conservation of solute concentration at each timestep, which is the desired behavior as we endeavor to model the ventricles as a closed system. Note that the total concentration would still increase in time, because we impose the concentration profile in the photoconversion region Ω_pc_ (region of interest 1).

Using the introduced approaches of handling the in- and outflow boundaries, the resulting abstract weak formulation for the cardiac-induced/no-cilia model is: find *c* ∈ *W* such that

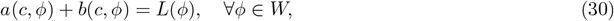

with the definitions

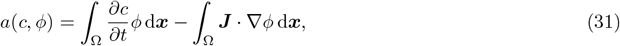

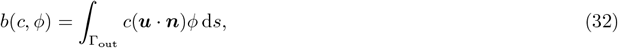

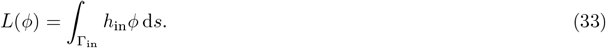

#### Baseline model

Since both free-slip and cilia tangential traction boundaries have the same no-flux boundary condition in the advection-diffusion equation, the weak formulation of the advection-diffusion equation for the baseline model is identical to that of the cardiac-induced/no-cilia model, defined by Equations (30), (32) and (33).

### A.4 Discretization of the advection-diffusion equation

We outline the discretization of the advection-diffusion weak formulation for the baseline model, defined by Equations (30), (32) and (33). The weak formulation is discretized with a symmetric interior-penalty method using discontinuous Galerkin finite elements.^[110]^ Exchanging the variables in Equation (30) with their discrete counterparts yields the discrete weak form: find *c*_*h*_ ∈ *W*_*h*_ such that

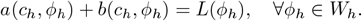

The space *W*_*h*_ is constructed with finite elements with discontinuous Lagrange polynomial basis functions:

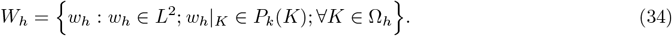

These elements allow for discontinuities in the discrete solution across element edges. To ensure a stable discretization with these elements, we add the stabilization

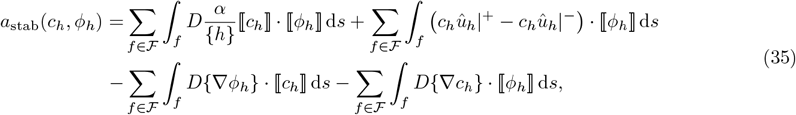

to the discrete weak formulation, where

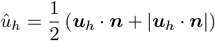

is used to upwind the velocity ***u***_*h*_ with respect to a facet. The stabilization parameter *α >* 0 must be chosen large enough to achieve stability. In Equation (35), the first two terms penalize jumps in the concentration flux across interior facets, and are needed to preserve coercivity of the bilinear form. The last two terms in *a*_stab_ are added to preserve symmetry and consistency of the bilinear form, respectively. The average and jump operators used in Equation (35) are defined, respectively, as

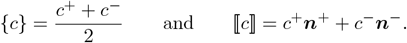

By including the stabilization *a*_stab_(*c*_*h*_, *ϕ*_*h*_), the final semi-discrete weak formulation of the advection-diffusion equation is: find *c*_*h*_ ∈ *W*_*h*_, such that

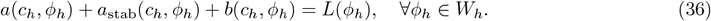

In time, we discretized Equation (36) with the second-order backward difference scheme BDF2.^[111]^ This scheme approximates the time derivative as

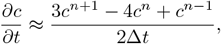

where *n* denotes the timestep index. All of the other terms in Equation (36) are evaluated at timestep *n* + 1. The method is implicit and of second order in time. We solved the fully discrete weak formulation of the advection-diffusion equation with the penalty parameter value *α* = 50.

## B Verification

### B.1 Method of Manufactured Solutions

Let 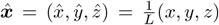 be the spatial coordinates scaled by the length *L*, which is the length of the computational domain Ω in the *x*-direction. Letting ***ϕ*** be the vector-valued function

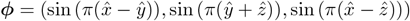

and defining the velocity as

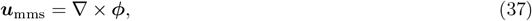

we have by construction that the velocity is divergence-free, owing to the identity∇· (∇ × ***F***) = 0 for an arbitrary vector field ***F***. The pressure was defined as

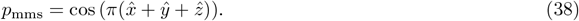

Using the Method of Manufactured Solutions,^[120]^ we construct a problem with the Stokes equations where Equation (37) and Equation (38) are the solutions. To this end, we consider the general form of the steady Stokes equations

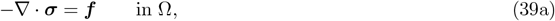

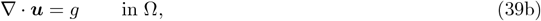

with ***σ***(***u***, *p*) = 2*µ****ε***(***u***) − *p***I** and 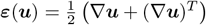, and the boundary condition

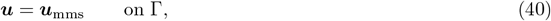

where Γ = *∂*Ω is the boundary of the domain. Since the velocity field defined by Equation (37) is divergence-free by construction, *g* = 0. Setting the external force ***f*** = −∇ · ***σ***(***u***_mms_, *p*_mms_) using Equation (37) and Equation (38), the solutions ***u*** and *p* to Equation (39) will be the ones defined by Equation (37) and Equation (38), as long as we enforce equality in the mean pressures, i.e. ⎰_Ω_(*p*_*h*_ − *p*_mms_) d***x*** = 0. Convergence rates are then readily calculated by comparing the numerical approximations of ***u*** and *p* with ***u***_mms_ and *p*_mms_ on meshes of varying grid resolution.

### B.2 orms and convergence order

Errors of the velocity ***u***_*h*_ and pressure *p*_*h*_ approximated with finite elements were measured in **H**^1^ and *L*^2^ error norms, respectively. With the exact velocity field ***u***_ex_, the **H**^1^ error semi-norm was calculated as

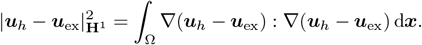

The *L*^2^ error norms for the pressure were calculated as

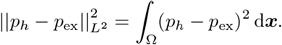

The approximated functions were interpolated onto elements in a space of piecewise-continuous Lagrange polynomials of three polynomial degrees higher than the elements used to approximate the functions, meaning that polynomial degrees 4 and 3 were used for the velocity and pressure errors, respectively. Numerical integration was performed with a Gauss quadrature rule, with the quadrature rule degree equal to the polynomial degree of the integrated finite element functions.

Convergence rates reported in Appendix B.3 were approximated in the following way. Define the error *e*_*k*_ for a given mesh resolution *h*_*k*_. Based on the assumption that the error scales as *e*_*k*_ ≈ *Ch*_*k*_ ^*η*^, where *C* is a constant independent of *h*_*k*_, the convergence order *η* was approximated as

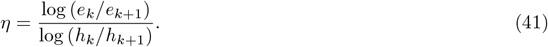

We define the approximation of the infinity norm of a finite element function *ϕ* = *ϕ*_*i*_*ψ*_*i*_(***x***) as

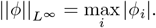

Here *ψ*_*i*_ are the basis functions and *ϕ*_*i*_ are the degrees of freedom defined as point evaluations.

### B.3 Numerical verification of model implementation

A cylinder geometry meshed with varying degrees of refinement was used to perform convergence analysis of the numerical method used to solve the Stokes equations. The cylinder had unit length and a diameter of one half length unit. For each mesh refinement, the edge lengths were approximately halved. Since the refinement was not uniform, we used the ratio of the minimal edge lengths *h*_min,*k*_*/h*_min,*k*+1_ between two successive refinements when calculating the convergence rate using Equation (41). For the analysis, we used tangential traction boundary conditions on all of the cylinder surface. Using the Method of Manufactured Solutions^[120]^ with the velocity and pressure expressions defined in Appendix B.1 for ***u***_ex_ and *p*_ex_, we calculated the error of the velocity approximation ***u***_*h*_ in the **L**^2^ norm and **H**^1^ semi-norm, and the error of the pressure approximation *p*_*h*_ in the *L*^2^ norm (Table 3). Based on the error calculations, we calculated the experimental order of convergence, as defined in Appendix B.2. We observe convergence in all of the reported error norms. Furthermore, we report the error in calculation of the maximum velocities, and observe convergence in this error as well. Finally, we calculated the *L*^2^ norm of the divergence ∇ · ***u***_*h*_ to verify the property of mass conservation.

**Table 3:**
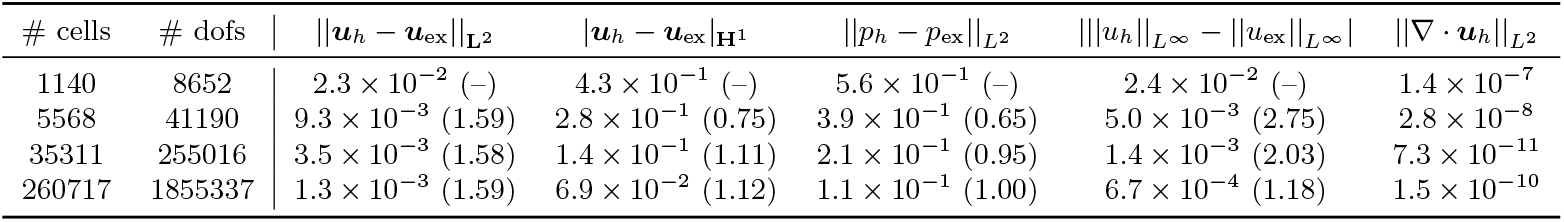
Error norms for the convergence study performed with a cylinder mesh with the cilia-driven/no-cardiac flow model using *µ* = 1. For all error norms, convergence rates calculated with Equation (41) are given in parentheses.

| # cells | # dofs | $\ \mathbf{u}_h - \mathbf{u}_{\text{ex}}\ _{\mathbf{L}^2}$ | $\ \mathbf{u}_h - \mathbf{u}_{\text{ex}}\ _{\mathbf{H}^1}$ | $\ p_h - p_{\text{ex}}\ _{L^2}$ | $\ u_h\ _{L^\infty} - \ u_{\text{ex}}\ _{L^\infty}$ | $\ \nabla \cdot \mathbf{u}_h\ _{L^2}$ |
| --- | --- | --- | --- | --- | --- | --- |
| 1140 | 8652 | $2.3 \times 10^{-2}$ (-) | $4.3 \times 10^{-1}$ (-) | $5.6 \times 10^{-1}$ (-) | $2.4 \times 10^{-2}$ (-) | $1.4 \times 10^{-7}$ |
| 5568 | 41190 | $9.3 \times 10^{-3}$ (1.59) | $2.8 \times 10^{-1}$ (0.75) | $3.9 \times 10^{-1}$ (0.65) | $5.0 \times 10^{-3}$ (2.75) | $2.8 \times 10^{-8}$ |
| 35311 | 255016 | $3.5 \times 10^{-3}$ (1.58) | $1.4 \times 10^{-1}$ (1.11) | $2.1 \times 10^{-1}$ (0.95) | $1.4 \times 10^{-3}$ (2.03) | $7.3 \times 10^{-11}$ |
| 260717 | 1855337 | $1.3 \times 10^{-3}$ (1.59) | $6.9 \times 10^{-2}$ (1.12) | $1.1 \times 10^{-1}$ (1.00) | $6.7 \times 10^{-4}$ (1.18) | $1.5 \times 10^{-10}$ |

To verify the model implementation with the zebrafish brain ventricles mesh, we compared velocity norms for varying degrees of refinement of the brain ventricles mesh (Table 4). We used the original mesh obtained by meshing the surface geometry using fTetWild;^[96]^ the region around the photoconversion region is not refined on the mesh used for the convergence study. In the numerical experiments, we used the cilia traction vector ***τ*** = *τλ*(***x***)*P* _*n*_ (***r***) defined in Table 2 and *γ* = 10 for the BDM penalty parameter.

**Table 4:** Velocity norms under mesh refinement of the brain ventricles mesh using the cilia-driven/no-cardiac flow model.

| # cells | # dofs | $\ u_h\ _{L^\infty} \left[ \frac{\text{mm}}{\text{s}} \right]$ | $\frac{1}{F} \ u_h\ _{L^\infty} \left[ \frac{1}{\text{Pa} \cdot \text{mm} \cdot \text{s}} \right]$ | $\frac{1}{ \Omega } \ \mathbf{u}_h\ _{\mathbf{L}^2} \left[ \frac{\text{mm}}{\text{s}} \right]$ | $\ \nabla \cdot \mathbf{u}_h\ _{L^2} \left[ \frac{\text{mm}^3}{\text{s}^2} \right]$ |
| --- | --- | --- | --- | --- | --- |
| 132134 | 943820 | 0.026 | $3.8 \times 10^3$ | $2.2 \times 10^{-3}$ | $1.9 \times 10^{-16}$ |
| 180901 | 1289881 | 0.027 | $3.7 \times 10^3$ | $2.2 \times 10^{-3}$ | $3.0 \times 10^{-16}$ |
| 262598 | 1870703 | 0.027 | $3.7 \times 10^3$ | $2.3 \times 10^{-3}$ | $6.7 \times 10^{-16}$ |
| 395339 | 2811668 | 0.028 | $3.6 \times 10^3$ | $2.2 \times 10^{-3}$ | $2.0 \times 10^{-14}$ |
| 627439 | 4448761 | 0.027 | $3.5 \times 10^3$ | $2.2 \times 10^{-3}$ | $4.9 \times 10^{-14}$ |

For all meshes, we report the maximum discrete velocity magnitude 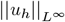, with 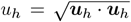. We also report scaled by the total force 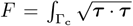 d*s* applied on the cilia boundary Γ_c_. We chose this measure using *F* as a scale because the regions of the mesh marked as cilia-populated change slightly with every mesh refinement, owing to the unstructured organization of the mesh cells. These changes in the surface area over which the tangential traction is applied result in changes to 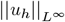, since the maximum velocity magnitude is a result of the total amount of force *F* applied on the cilia boundary.

To account for the fact that the volume of the domain |Ω| = _Ω_ d***x*** also changes slightly with mesh refinement, we report the **L**^2^ norm of ***u***_*h*_ scaled by |Ω|. Finally, we provide verification of mass conservation by reporting the *L*^2^ norms of the divergence∇ · ***u***_*h*_. We remark that the units used in the model implementation was millimeters, which is why Table 4 reports values in units of millimeters, instead of micrometers which has been used elsewhere in the paper.

### B.4 Discussion of the numerical approximation and convergence rates

A priori error estimates for the Brezzi-Douglas-Marini and discontinuous Galerkin discretization scheme BDM_*k*_-DG_*k*−1_ are naturally derived in a mesh-dependent broken **H**^1^-norm for ***u*** and the *L*^2^-norm for the pressure. The respective error norms are expected to decay with order *k*.^[121]^ For the sake of simplicity, Table 3 reports the velocity errors in the standard **H**^1^ (semi)norm. Using polynomial degree *k* = 1, we observe linear convergence in the velocity in the **H**^1^ error norm and the pressure in the *L*^2^ error norm on the cylinder mesh. For the ventricles meshes, we observe consistent values of the reported quantities, verifying the model implementation. The BDM_1_-DG_0_ scheme that we employ for the Stokes equations is computationally more expensive than more common approaches based on e.g. Taylor-Hood discretization^[82]^ (*P*_2_ − *P*_1_ elements) or lower-order enriched elements (e.g. MINI).^[83]^ We opted for the BDM-DG scheme because of its exactly-mass-conserving property, since numerical experiments proved that a divergence-free velocity field was fundamental to attain a stable discretization of the advection-diffusion equation in advection-dominated regions. First, we attempted to discretize the Stokes equations with Taylor-Hood elements, but with the resulting numerical approximation of the velocity ***u***_*h*_, we were unable to achieve stability of the advection-diffusion discretization. Numerical experiments identified non-zero divergence of ***u***_*h*_ as the root cause of these stability issues. On the standard computational mesh, the *L*^2^ norm of ∇·***u***_*h*_ was in the order of 10^−6^, while the maximum value of the divergence of the velocity was of order unity. On a refined mesh with roughly six times as many computational cells as the standard mesh, the numerical values of 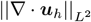 and 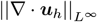 remained practically unchanged. This is a result of the fact that incompressibility is only achieved in a weak sense when employing Taylor-Hood elements. The advective part of the total flux is

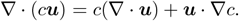

In the case that ∇ ·***u*** ≠ 0, a source (or sink) term for the concentration is introduced. This may introduce errors in the numerical approximation of *c* and lead to instabilities and spurious oscillations, especially in the case of advection-dominated transport.^[84–86]^ The BDM_1_-DG_0_ scheme avoids these issues, since ∇ ·***u***_*h*_ = 0 is satisfied exactly as a result of the element definitions.

In addition to its favorable mass-conserving property, the BDM-DG scheme lets us impose the impermeability condition strongly without any tailored approach, since this boundary condition appears naturally in the weak form of the Stokes equations (Appendix A.1) and the BDM elements control the normal component of the velocity. If instead (for example) Taylor-Hood elements were to be used, the impermeability boundary condition must be handled with for example a Nitsche method^[116]^ or with a Lagrange multiplier.^[122, 123]^

## C Quantification of velocities in particle tracking

Particle tracking data was analyzed.^[14]^ For four zebrafish, five recordings with a few particles for each fish were carried out. The particles tracked were in the duct between the rhombencephalic ventricle and the diencephalic ventricle. Frequency of image acquisition was 20.22 Hz. The collected data constituted *x* (rostrocaudal axis) and *z* (dorsoventral axis) positions of the particles at every time instant of image acquisition. The velocities *u*_*x*_ and *u*_*z*_ were determined by calculating the change in positions Δ*x* and Δ*z* relative to the change in time Δ*t* between two subsequent images, Δ*t* being equal to the frequency of acquisition. Mean velocities *ū*_*x*_ (Figure 7a) and *ū*_*z*_ (Figure 7b), which serve as a quantification of particle drift, were calculated by taking the difference between the last and first positions of a particle.

**Figure 7:**
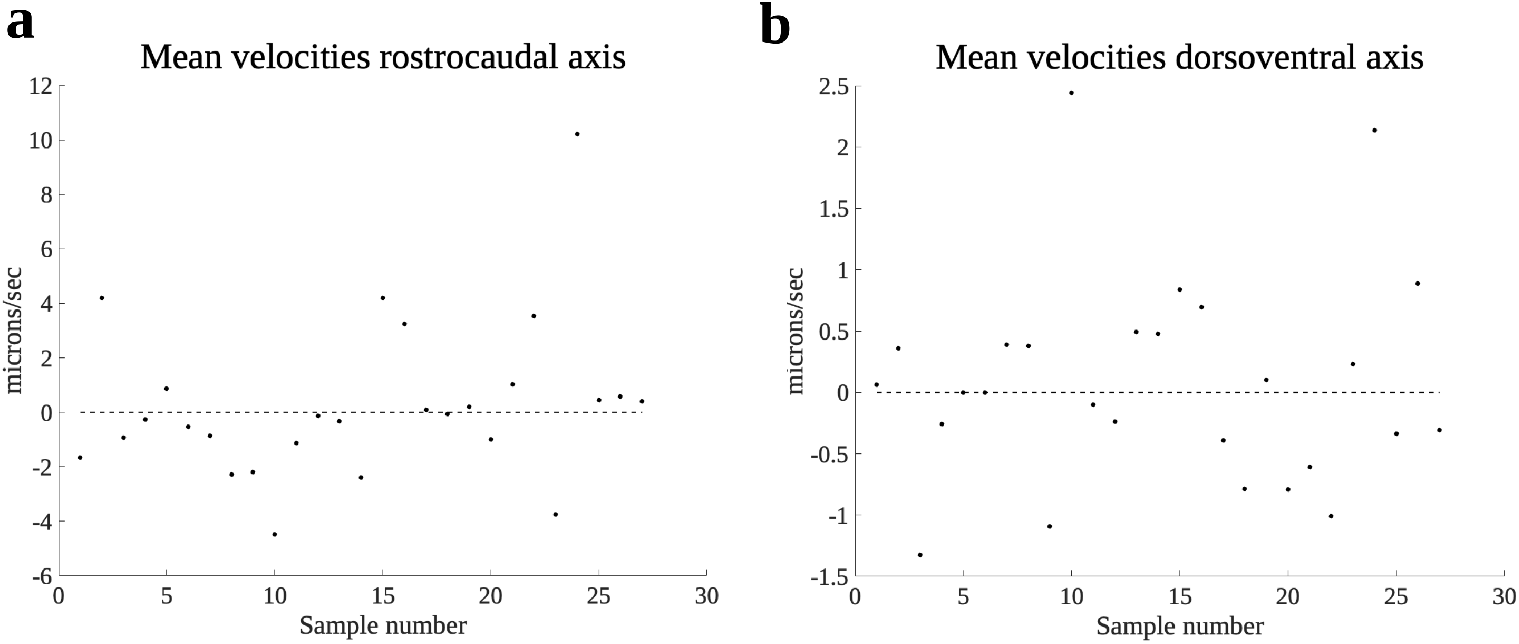
Particle tracking data. Mean velocities in along the rostrocaudal axis (*x* axis, **a**) and along the dorsoventral axis (*z* axis, **b**) directions of the particles tracked. The dashed horizontal lines designate zero.

Particle movement varied over the different measurements. The mean of *ū*_*x*_ was 0.26 µm/s and the mean of *ū*_*z*_ was 0.08 µm/s. The maximum mean velocities in absolute values were 10.2 µm/s and 2.4 µm/s for *ū*_*x*_ and *ū*_*z*_, respectively. We used the cardiac-induced/no-cilia flow model to tune the pressure boundary condition amplitude *A*. The choice of *A* = 1.5 mPa was used since: (i) it yields a maximum velocity magnitude of 4.8 µm/s, which is close to the maximum observed in the particle tracking data when rejecting the outlier at around 10 µm/s; (ii) it yields a time-averaged mean velocity of 0.89 µm/s, which is close to the mean observed in the particle tracking data.

## Footnotes

1 To simplify the exposition we focus only on spatial regularity. In particular, ***v*** ∈ **H**^1^(Ω) is understood as ***v***(·, *t*) ∈ **H**^**1**^ (Ω) for all *t >* 0. Similar convention is applied also when discussing the advection-diffusion equation in Appendix A.3.

## Notes

https://doi.org/10.5281/zenodo.15194757

### Competing Interest Statement

The authors have declared no competing interest.

