## Supplemental information for "Advection versus diffusion in brain ventricular transport"

**Video S1:** CSF flow simulation with motile cilia and cardiac pulsatility. Initial view is lateral (xz-plane). Velocity vectors are scaled by magnitude, and colored by the magnitude signed with the x component, such that rostrocaudal flow is red and caudorostral flow is blue. The video spans 24 cardiac cycles at 22 fps rate (half of real-time).

**Video S2:** Transport of secreted and photoconverted Dendra2 in the zebrafish ventricle. Dual color time-lapse confocal images show the location of unconverted (green) and converted (magenta) signals over time. Acquisition rate: 2.67s per frame. Photoconversion is performed from frame 11 to the end of the recording (frame 300).
